# Dimensional and Categorical Predictors of Theory of Mind Network Activation: A transdiagnostic fMRI Study in Psychiatric Inpatients

**DOI:** 10.64898/2026.09.21.753139

**Authors:** Pauline Maria Nagel, Simon Hefner, Sarah Said-Yürekli, Daniel Freinhofer, Kornelius Winds, Wolfgang Aichhorn, Martin Kronbichler, Lisa Kronbichler

## Abstract

Socio-cognitive alterations are widely observed across different psychiatric conditions, yet it remains unclear whether these deficits map onto distinct diagnostic categories or represent a shared transdiagnostic phenotype, driven by dimensional psychopathological symptom severity.This study evaluates Theory of Mind (ToM) network activation during a naturalistic functional magnetic resonance imaging (fMRI) movie paradigm in 313 participants, including 91 healthy controls and 222 psychiatric inpatients across five distinct diagnostic clusters (substance use, schizophrenia spectrum, affective, anxiety, and personality disorders).

Frequentist analyses revealed decreased ToM-selective activity in psychiatric patients, compared to controls across all ToM regions of interest (ROIs). No significant difference emerged between diagnostic groups, but ROI activation correlated negatively with Adult Self-Report (ASR) scales (*Total Problems, Internalising, Externalising*). Subsequent Bayesian analysis of covariance (ANCOVAs) consistently favoured models that excluded traditional categorical diagnoses, suggesting that dimensional measures, especially Total Problems and Internalising, provided substantially greater predictive utility for ToM-related regional brain activation. Our findings challenge traditional categorical classifications in neurobiological research as they suggest that atypical mentalising might reflect a shared transdiagnostic vulnerability.

**Highlights:**

- Reduced Theory of Mind specific activation in psychiatric inpatients presents as a shared, transdiagnostic feature.
- Theory of Mind specific activation shows no disorder-specific patterns across five distinct clinical diagnostic clusters.
- Bayesian analyses consistently favour continuous psychopathology dimensions over categorical classifications in predicting Theory of Mind specific activation

## 1. Introduction

Social cognition, particularly the ability to attribute mental states to others (Theory of Mind; ToM), is a fundamental prerequisite for successful interpersonal functioning (Arioli & Canessa, 2019; Van Overwalle, 2009). Disruptions in these mentalising processes, alongside altered activation in the underlying cortical network (comprising, e.g., the medial prefrontal cortex, temporoparietal junction, and precuneus), are widely recognised as hallmarks of severe psychiatric illness (Cotter et al., 2018a; Zhu et al., 2026).

Historically, neuroimaging research has approached these deficits through a categorical lens, seeking to identify disorder-specific neural signatures by comparing isolated clinical populations, such as schizophrenia or major depressive disorder (MDD), against healthy controls (Cusi et al., 2012; Kronbichler et al., 2017; Segal et al., 2023). This approach has generated a wealth of literature suggesting divergent, diagnosis-specific activation patterns across the mentalising network (Malhi et al., 2008; Porcelli et al., 2019). To illustrate, studies identified frontoparietal hypo- and hyper-activation across the medial prefrontal cortex (mPFC) and the temporoparietal junction (TPJ) in schizophrenia (Kronbichler et al., 2017), attenuated prefrontal engagement coupled with enhanced limbic activity in affective disorders like MDD (Cusi et al., 2012; Porcelli et al., 2019), disruption in superior temporal sulcus in anxiety disorders (Cui et al., 2017) and altered amygdala and medial frontal activation in personality disorder (Frick et al., 2012).

Such categorical thinking is increasingly challenged by substantial methodological and conceptual limitations (Cicero et al., 2024; Cotter et al., 2018a; Hengartner & Lehmann, 2017; Kotov et al., 2017). Disentangling disorder-specific cortical activation patterns remains highly problematic, as the prevailing literature relies on heterogeneous ToM tasks with divergent cortical activation profiles (Schurz et al., 2014) applied to single-patient cohorts (Van Neerven et al., 2021). Crucially, direct transdiagnostic comparisons across multiple psychiatric groups remain rare (Fusar-Poli et al., 2019; Oliver et al., 2024). As a result, it remains an open question whether the identified neural ToM profiles for specific psychiatric illnesses reflect genuine categorical boundaries, or simply the methodological noise of disparate study designs (Tsui et al., 2025).

Furthermore, the validity of traditional nosological boundaries (e.g., ICD or DSM categories) in capturing underlying biological and functional realities is undergoing a critical re-evaluation (Farías Venegas et al., 2025; Fusar-Poli et al., 2019; Leucht et al., 2024) and recent empirical evidence strongly suggests that dimensional models of psychopathology offer significantly greater explanatory power than categorical diagnoses (Caspi & Moffitt, 2018; Nagel et al., 2026). For instance, transdiagnostic dimensions, particularly broad general psychopathology and domains of dysregulation, have been shown to significantly outperform traditional categorical diagnoses in predicting real-world functional disability and social outcomes (Braak et al., 2022; Nagel et al., 2026). Converging evidence from psychiatric genetics similarly highlights the superiority of dimensional over categorical approaches (Mallard et al., 2022; Sprooten et al., 2022; Thomas, 2026; Waszczuk et al., 2020), demonstrating that broad transdiagnostic dimensions, such as the general p-factor, capture an extensive transdiagnostic architecture across clinical phenotypes. This suggests that the field’s historical reliance on discrete diagnoses may partly explain the limited translational progress in psychiatric neuroscience over recent decades. Despite this paradigm shift in clinical research, neurobiological studies directly contrasting categorical and dimensional predictors within a unified, multi-group design remain scarce (Oliver et al., 2024).

To address the limitations of isolated assessments, the present study employed a naturalistic fMRI ToM paradigm (movie viewing; Jacoby et al., 2016; Richardson et al., 2018). We investigated a large, transdiagnostic inpatient sample encompassing five distinct clinical clusters (substance use, schizophrenia spectrum, affective, anxiety, and personality disorders) alongside healthy controls.

By doing so, we aimed to achieve the following objectives: First, we examined whether the viewing paradigm used in this study reveals altered activation patterns for psychiatric patients (all groups combined) compared to healthy controls in areas associated with ToM. Second, we directly contrasted ToM network activation between the five diagnostic clusters. We reasoned that if traditional diagnostic classifications relate to distinct ToM-specific altered activation patterns, disorder-specific differences in the examined ROIs would emerge (Miller et al., 2023). Third, we examined whether dimensional measures more closely map onto cortical activation by 1) examining the existence and direction of correlations between measures of psychopathology and blood-oxygen-level-dependent (BOLD) response and 2) comparing the predictive utility of the categorical diagnoses against dimensional psychopathology with respect to BOLD response by means of Bayesian model comparisons (Parkes et al., 2021; Voldsbekk et al., 2023). In doing so, we aim to determine whether atypical ToM activations are driven by specific categorical diagnoses, or rather reflect a shared, dimensional vulnerability across the psychiatric spectrum.

## 2. Methods

### 2.1 Participants

This study analysed data from 313 participants, including 91 healthy controls, who were recruited from the University of Salzburg. 222 patients were recruited from the inpatient units of the Department of Psychiatry, Psychotherapy, and Psychosomatics at the Christian Doppler Clinic in Salzburg between July 2018 and March 2025. Specifically, 24 patients had a substance use disorder (SUD), 44 had a schizophrenia spectrum disorder (SZD), 102 had an affective disorder (AD), 32 had an anxiety disorder (ANX), and 20 had a personality disorder (PD). All patients were clinically stable and had received a formal ICD-10 diagnosis, which was confirmed before participation by certified psychiatrists or clinical psychologists using the Structured Clinical Interview for DSM Disorders (SCID) (Shabani et al., 2021). Patient availability determined the sample size rather than an a priori power analysis.

Exclusion criteria for all participants included anamnestic neurological injuries or impairments (e.g., severe head trauma). Recruited healthy controls were screened for mental and physical health. Only individuals able to give informed consent (age > 18 years, not under external legal custody) were eligible for inclusion. The study was approved by the local ethics committee (Ethikkommission für das Bundesland Salzburg), and all methods were performed in accordance with their relevant guidelines and regulations and in accordance with the Declaration of Helsinki.

Age and Gender distribution are presented in Table 1. Educational information for all participants is provided in Supplementary Section 1.

**Table 1.** Demographics.

| **Diagnosis** | **N** | **Female (%)** | **Male (%)** | **Mean Age (SD)** |
| --- | --- | --- | --- | --- |
| **SUD (F10-F19)** | 24 | 3 (12,5%) | 21 (87,5%) | 39,04 (10,61) |
| **SZD (F20-F29)** | 44 | 16 (36,4%) | 28 (63,6%) | 36,50 (11,87) |
| **AD (F30-F39)** | 102 | 49 (48,0%) | 53 (52,0%) | 38,03 (15,07) |
| **ANX (F40-F49)** | 32 | 19 (59,4%) | 13 (40,6%) | 35,75 (13,57) |
| **PD (F60-F69)** | 20 | 14 (70,0%) | 6 (30,0%) | 24,50 (9,64) |
| **Healthy Controls** | 91 | 44 (48,4%) | 47 (51,6%) | 34,74 (14,72) |

### 2.2 Variables

#### 2.2.1 Behavioural Assessment

Dimensional psychopathology was quantified using the ASR (Achenbach & Dumenci, 2003)), where we obtained the scales *Internalising*, *Externalising,* and *Total Problems*.

#### 2.2.2 Natural Viewing Paradigm

To evaluate social cognition, participants watched a 5.49-minute muted version of the Disney-Pixar animated film Partly Cloudy in the scanner. Originally validated by Jacoby et al. (Jacoby et al., 2016), dynamic segments of the film isolate ToM processing (focusing on characters’ beliefs, motives and mental states) from baseline bodily sensations. Importantly, the functional dissociation between the ToM and Pain networks elicited by this type of naturalistic paradigm has been demonstrated in children as young as three years, with network specialisation increasing throughout childhood (Richardson et al., 2018). In-scanner stimulus presentation was preceded by a 10-second fixation rest block. Participants were instructed to watch the movie attentively.

### 2.3 Neuroimaging Data Acquisition and Preprocessing

#### 2.3.1 fMRI Parameters

Functional imaging data were acquired with a Siemens Magnetom Prisma 3 Tesla scanner (Siemens AG, Erlangen, Germany) using a 64-channel head-coil. Functional images sensitive to BOLD contrast were acquired with a T2* weighted gradient echo EPI (echo-planar imaging) sequence (Repetition Time [TR] = 1050 ms, Echo Time [TE] = 32.0 ms, matrix = 80 × 80, Field of View [FOV] = 192 mm, flip angle = 45°). Fifty-six slices with a slice thickness of 2.4 mm and no slice gap were acquired within the TR. The functional run consisted of 354 total volumes, beginning with 6 dummy scans, which were later discarded. A gradient echo field map (TR 623.0 ms, TE 1 = 4.92 ms, TE 2 = 7.38 ms) and a high-resolution (0.8 × 0.8 × 0.8 mm) structural scan with a T1-weighted MP-RAGE (magnetization-prepared rapid gradient-echo) sequence were acquired from each participant.

For preprocessing and statistical analysis, SPM12 software (http://www.fil.ion.ucl.ac.uk/spm/), running in a MATLAB R2013a environment (The MathWorks Inc., 2013), and additional functions from AFNI (https://afni.nimh.nih.gov/) were used. Further details on fMRI preprocessing are provided in the Supplementary Section 2.

Contrast images for the effects of interest were calculated at the first level, using the timings (onset and length) for Theory of Mind and Pain events provided by Richardson et al. (2018) (for the exact timings, see Richardson’s Supplementary Table 3). Effects were rescaled to increase statistical sensitivity and decrease inter-individual variability by the Vascular auto-rescaling of fMRI (VasA fMRI) technique (Kazan et al., 2016).

For the ROI analyses, we created 7mm spherical masks around the center coordinates of the ToM network regions as described in Richardson et al. (2018) including right TPJ (*x* = 48 ; *y* = −60 ; *z* = 30), left TPJ (*x* = −48; *y* = −62 ; *z* = 30), Precuneus (PC; *x* = 0; *y* = −54; *z* = 34), Dorsomedial Prefrontal Cortex (DMPFC; *x* = −6; *y* = 54; *z* = 36), Ventromedial Prefrontal Cortex (VMPFC; *x* = −4; *y* = 56; *z* = − 16;) and Middle Medial Prefrontal Cortex (MMPFC; *x* = −4; *y* = 58; *z* = 16). We then extracted contrast estimates from the *ToM* > *Pain* first-level rescaled contrast image of each participant, averaging across all voxels within each ROI.

Subsequent analyses were done using JASP (*Jasp Team*, 2025).

### 2.4 Analysis of Covariance (ANCOVA)

To evaluate categorical differences in neural activation while controlling for age and sex, we performed two-tailed ANCOVAs. First, separate univariate ANCOVAs were conducted for each ToM ROI to evaluate differences in overall diagnostic clusters (combined clinical sample vs healthy controls), thereby establishing whether ToM-related neural alterations were present in this paradigm (Results section 3.1). Second, a Disorder-Specific Evaluation was performed to examine differences in activation patterns between patient samples. Univariate ANCOVAs were conducted for each ROI to systematically compare the five distinct diagnostic groups (SUD, SZD, AD, ANX, PD).

#### 2.4.1 Correlation Analyses

To examine the relationship between regional neural activation (beta estimates) and dimensional symptom severity, we performed bivariate correlation analyses across all investigated ToM ROIs. Specifically, we correlated the neural activation metrics with the three dimensional scales of the ASR: *Total Problems, Externalising*, and *Internalising*.

#### 2.4.2 Bayesian ANCOVAs

To evaluate whether dimensional symptom measures or categorical diagnosis yield stronger evidence for regional brain activation across our ROIs, we performed a series of Bayesian ANCOVA models. The dimensional measures were derived from the Adult Self-Report (ASR), specifically utilising the *Total Problems, Externalising, and Internalising* scales.

Due to substantial multicollinearity among the ASR scales, we conducted separate Bayesian ANCOVAs for each dimensional predictor rather than entering them into a single model. This approach ensures robust inference by avoiding arbitrary partitioning of shared variance. Models were compared against a standardised null model controlling for age and sex.

## 3. Results

### 3.1 Variance Analyses Patients vs. Controls

The individual ROI analyses consistently revealed decreased ToM-specific activation (i.e., a decreased ToM>Pain contrast) in patients compared with controls across all examined ToM ROIs (*F*s(1, 307) > 4.55, *p*s < .035). ANCOVA metrics, including post-hoc tests are listed in Supplementary Section 3.

Unsystematic regional variations emerged for the control predictors: A significant positive main effect of Sex was found for the LTPJ and the Precuneus (*F*s(1, 307) > 4.79, *p*s < .029), and a significant Sex and Group interaction emerged within the MMPFC (F(1, 307) = 5.30, p = .022). Age emerged as a significant negative covariate in the PC (F(1, 307) = 6.4, p = .012). No other regions showed significant associations with age.

### 3.2 Variance Analyses Between Patients

Between-patient samples ANCOVAs revealed no significant effect of Diagnosis cluster (*F*s(4, 209) < 1.27, *p*s > .28), indicating that brain metrics did not vary systematically across the five distinct diagnostic subgroups in either ROI. Furthermore, no significant main effects of Cluster*Sex interactions were observed (*F*s(4, 209) < 0.43, *p*s > 0.43). Activation patterns per group are shown in Figure 1, ANCOVA test statistics is provided in Supplementary Section 4.

**Figure 1.**
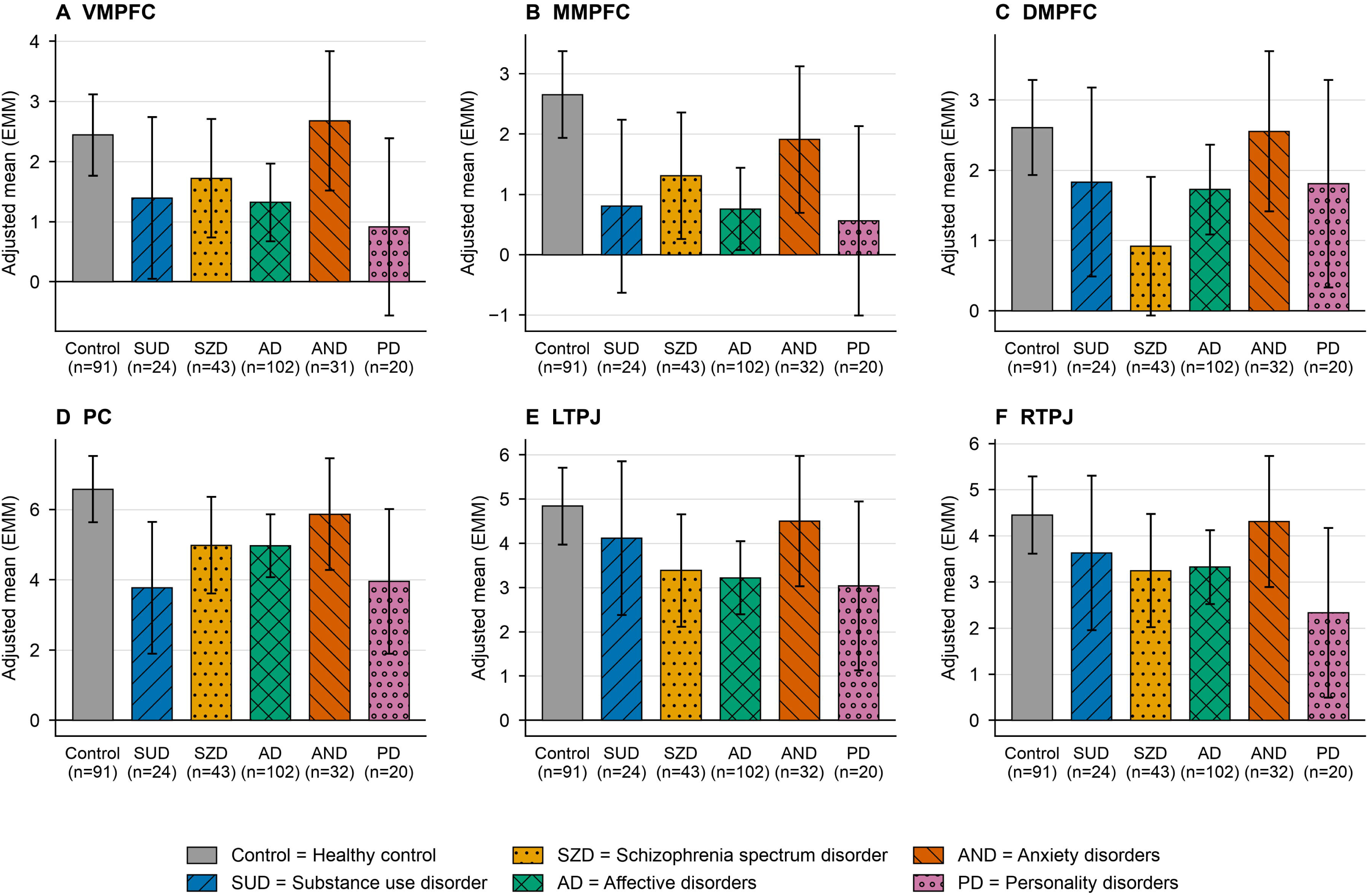
Age- and Sex-Adjusted ANCOVA Means by Region. Bar plots displaying the age- and sex-adjusted estimated marginal means (EMMeans) across ToM ROIs (VMPFC, MMPFC, DMPFC, PC, LTPJ, and RTPJ) for healthy controls and diagnostic clusters (SUD, SSD, AD, ANX, PD). Error bars represent 95% confidence intervals (CI).

Age was a significant negative predictor of activation in the DMPFC, PC, and RTPJ (*F*s(1, 209) > 4.57, *p*s < .033).

### 3.3 ROI and Symptom Correlations

Beta estimates across all regions of interest correlated negatively with all three ASR scales (see Figure 2 for a heatmap of all ROI-symptom correlations):

**Figure 2.**
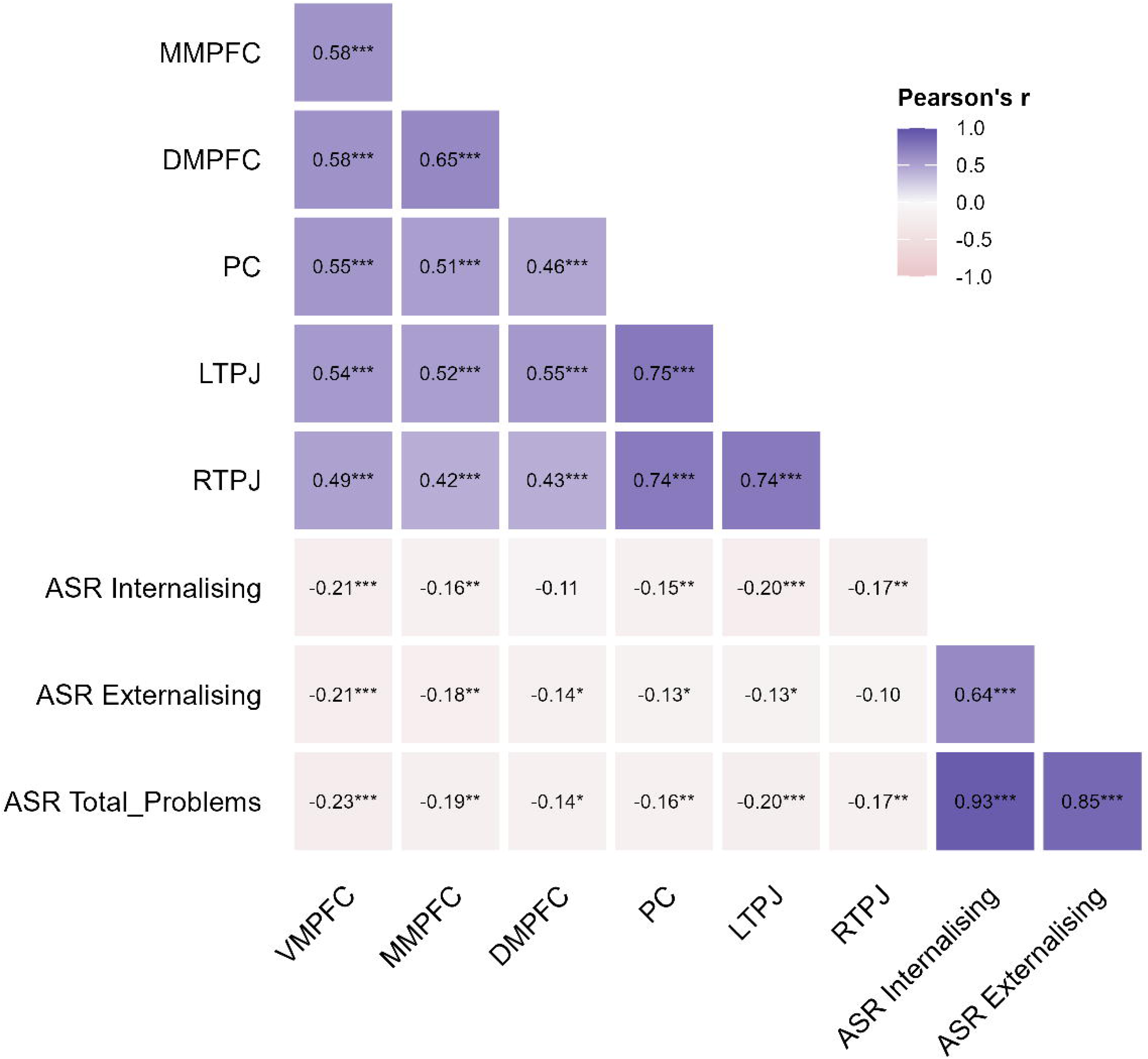
Pearson correlation heatmap of ROI measures and ASR behavioural scores. Displayed is the lower triangle of the correlation matrix. Colour scales range from pink (negative r) to purple (positive r). Asterisks indicate statistical significance: *p < .05, **p < .01, ***p < .001. VMPFC = ventromedial prefrontal cortex; MMPFC = medial mid-prefrontal cortex; DMPFC = dorsomedial prefrontal cortex; PC = precuneus; LTPJ/RTPJ = left/right temporoparietal junction; ASR = Adult Self-Report.

**Figure 3.**
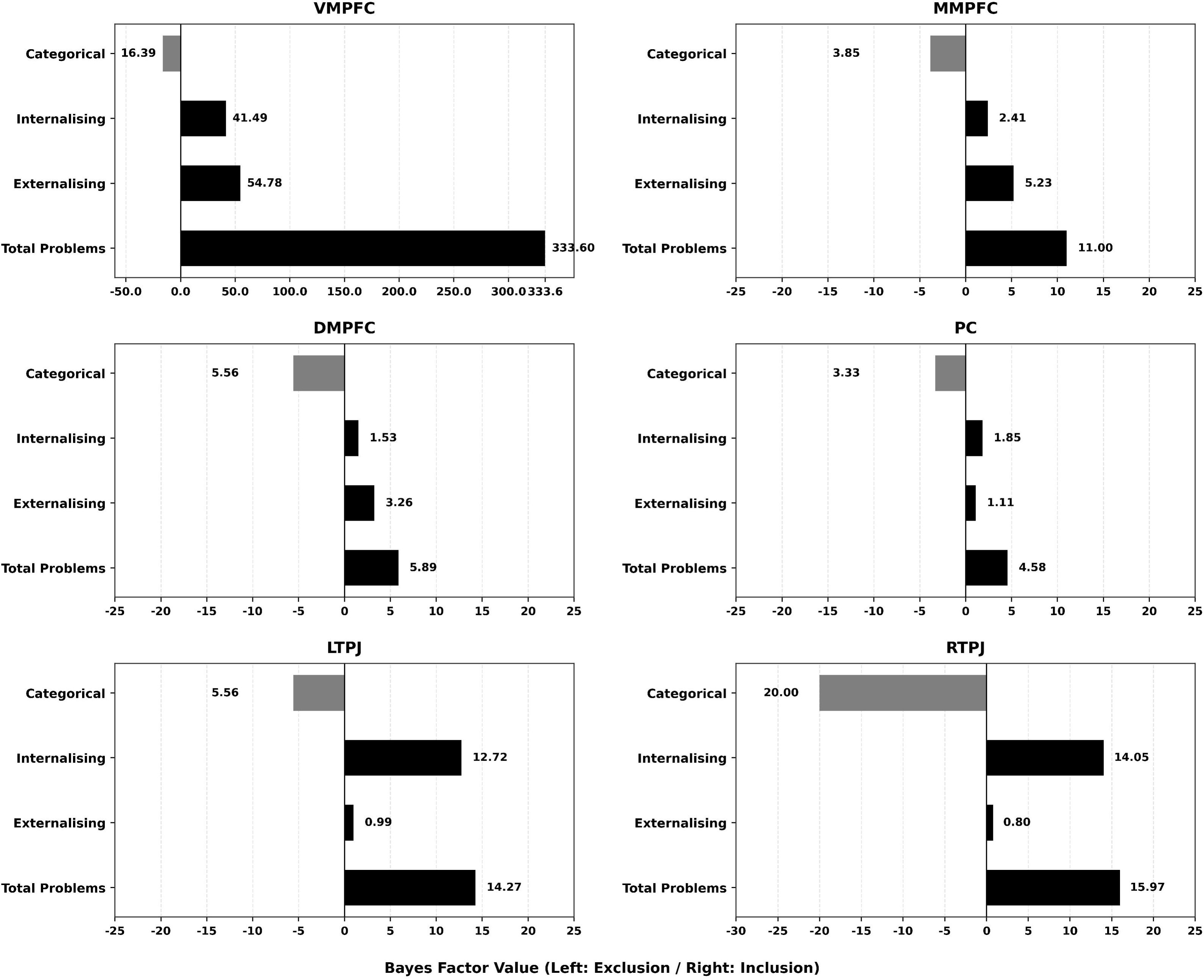
Log-transformed Bayes Factor (BF) strength across ToM ROIs. Positive values (right) indicate evidence for inclusion, while negative values (left) indicate evidence for exclusion. Note that the x-axis scale for the VMPFC subplot is independently adapted to accommodate the large positive Bayes factor for Total Problems (BF = 333.60), and the RTPJ subplot is scaled to clearly display the negative effect for Categorical predictors.

Higher general psychopathology (*ASR Total Problems*) was significantly associated with decreased neural activation across all investigated ROIs: VMPFC (*r = −.233, p < .001*), MMPFC (*r = −.187, p = .002*), DMPFC (*r = −.141, p = .018*), Precuneus (*r = −.159, p = .008*), LTPJ (*r = −.196, p < .001*), and RTPJ (*r = −.165, p = .005*).

Increased Internalising symptoms (*ASR Internalising*) were significantly linked to negative activation in the VMPFC (*r = −.207, p < .001*), LTPJ (*r = −.202, p < .001*) MMPFC (*r = −.164, p = .006*), RTPJ (*r = −.17, p = .004*), PC (*r = −.155, p = .009*). However, there was no significant correlation for the DMPFC (*r = −.0.11, p < .066*). Similarly, higher externalising (*ASR Externalising*) significantly correlated with lower activation in the VMPFC (*r = −.210, p < .001*), MMPFC (*r = −.185, p = .002*), DMPFC (*r = −.137, p = .022*), PC (*r = −.126, p = .034*), the LTPJ (*r = −.128, p = .32*), except for the RTPJ (*r = −.0.1, p = .104*).

### 3.4 Bayesian ANCOVAs

#### VMPFC

Regarding the VMPFC, *Total Problems* emerged as the strongest predictor (BFincl = 333.60), followed by *Externalising* (BF_incl_ = 54.78) and *Internalising* (BF_incl_ = 41.49). Evidence consistently favoured excluding the categorical diagnosis across all dimensional models (BF_excl_ ranging from 12.82 to 16.39; see Table 2 and Supplementary Sections 5.1, 6.1, and 7.1).

**Table 2.**
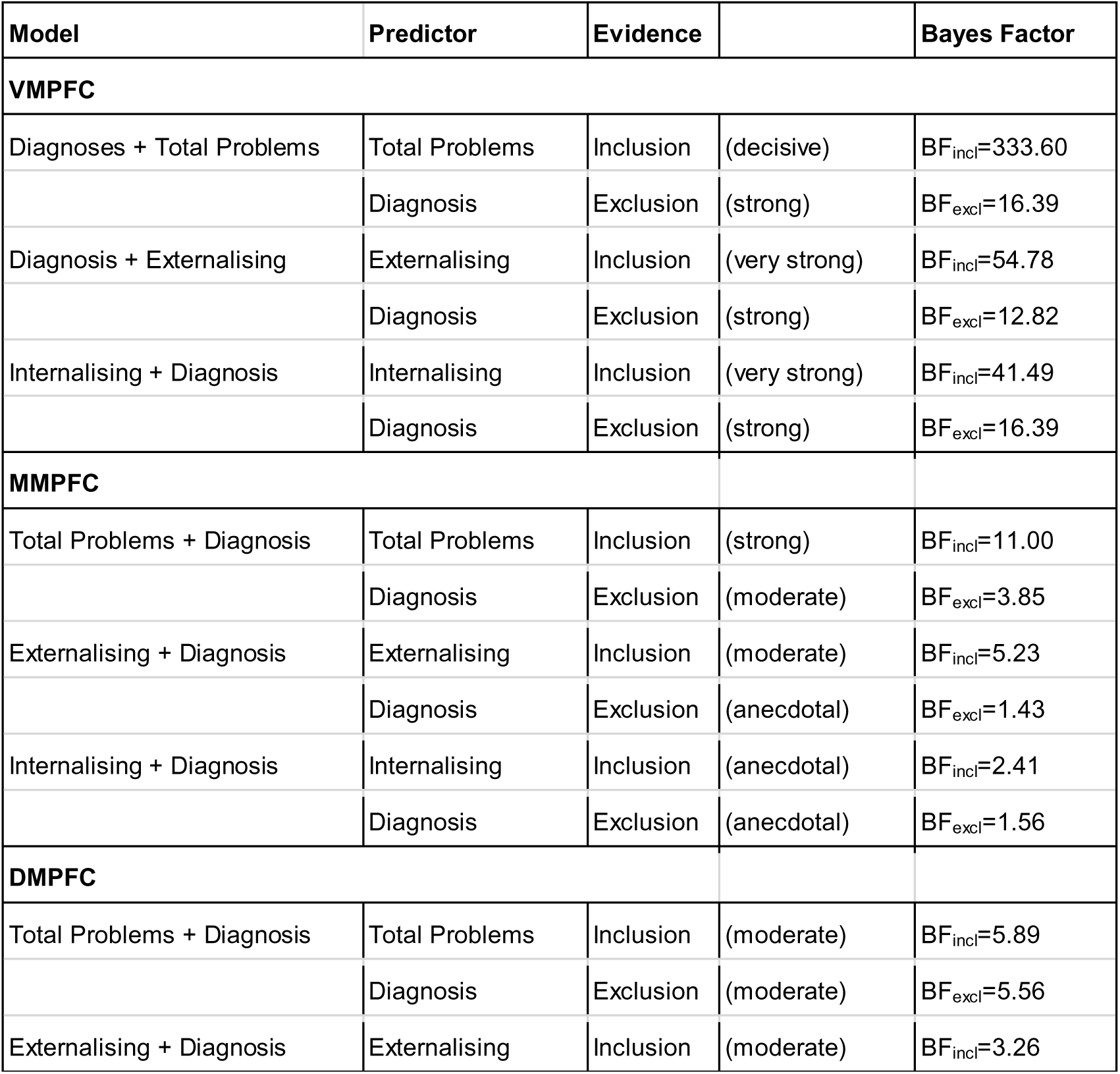
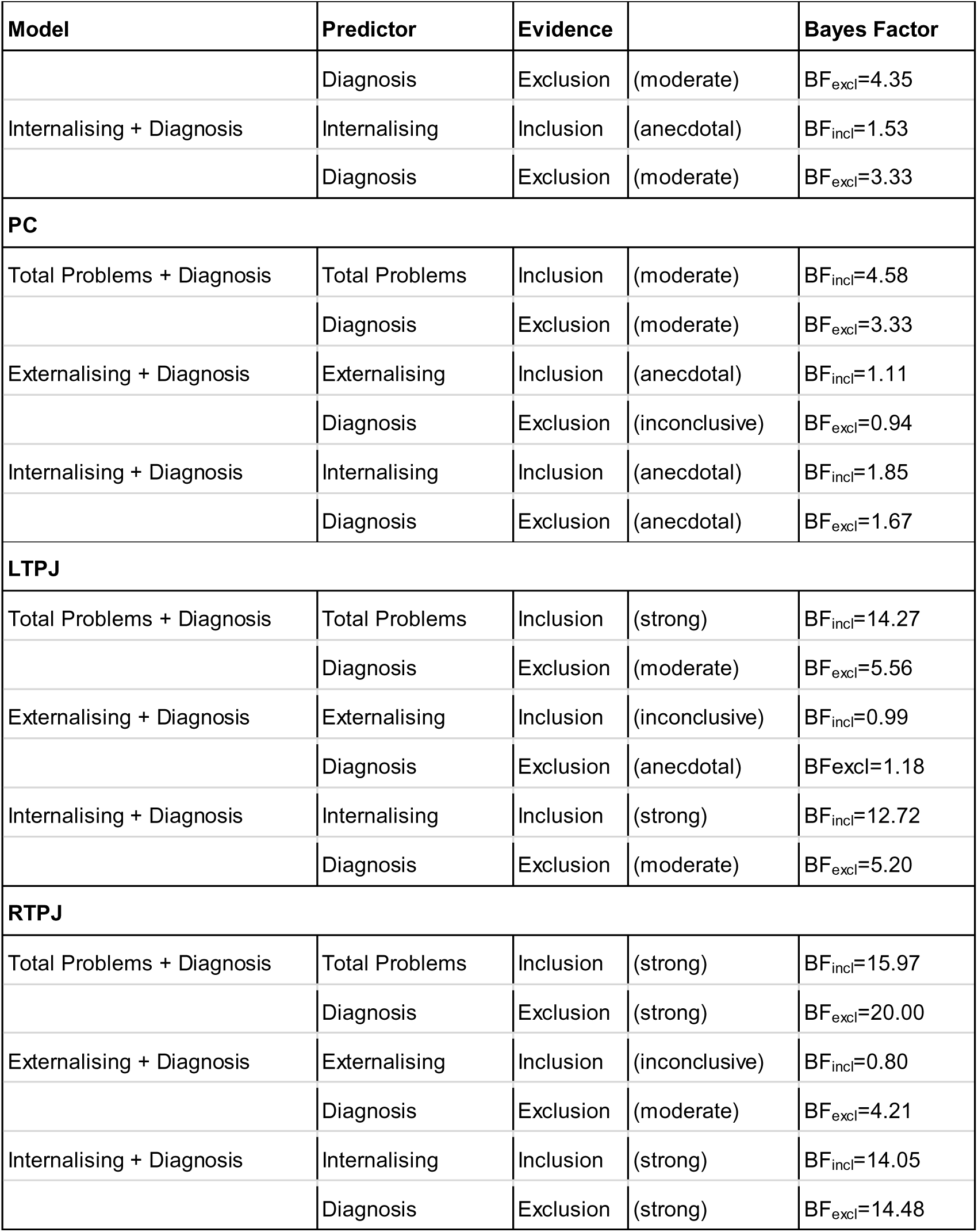
Summary of Bayesian ANCOVA results. Summary of Bayesian ANCOVA results evaluating the predictive utility of dimensional symptom measures (*Total Problems*, *Externalising*, *Internalising*) and the incremental value of categorical diagnosis across ROIs. Bayes factors for inclusion (BFincl) reflect evidence supporting dimensional predictors, whereas Bayes factors for exclusion (BFexcl) indicate evidence against adding the categorical diagnosis. Evidence categories follow Jeffreys’ classification criteria.

#### MMPFC

For the MMPFC, *Total Problems* showed strong evidence for inclusion (BF_incl_ = 11.00), *Externalising* moderate evidence (BF_incl_ = 5.23), and *Internalising* anecdotal evidence (BF_incl_ = 2.41). Categorical diagnoses consistently favoured exclusion (BFexcl = 1.43 to 3.85; see Table 2 and Supplementary Sections 5.2, 6.2 and 7.2 for details).

#### DMPFC

For the DMPFC, we found moderate evidence for including Total Problems (BF_incl_ = 5.89) and Externalising (BF_incl_ = 3.26), and anecdotal evidence for Internalising (BF_incl_ = 1.53). Categorical diagnosis consistently favoured moderate exclusion (BF_excl_ = 3.33 to 5.56; see Table 2 and Supplementary Sections 5.3, 6.3, and 7.3 for details).

#### PC

For the PC, *Total Problems* was the strongest predictor (BF_incl_ = 4.58), while *Externalising* (BF_incl_ = 1.11) and *Internalising* (BF_incl_ = 1.85) yielded anecdotal evidence. Categorical diagnoses favoured exclusion (BF_excl_ = 0.94 to 3.33; see Table 2 and Supplementary Sections 5.4, 6.4, and 7.4 for details). **LTPJ**

For the LTPJ, *Total Problems* (BF_incl_ = 14.27) and *Internalising* (BF_incl_ = 12.72) showed strong evidence for inclusion, whereas *Externalising* was inconclusive (BF_incl_ = 0.99). Categorical diagnoses favoured exclusion (BF_excl_ = 1.18 to 5.56; see Table 2 and Supplementary Sections 5.5, 6.5, and 7.6 for details).

#### RTPJ

This pattern repeated for the RTPJ, with *Total Problems* (BF_incl_ = 15.97) and *Internalising* (BF_incl_ = 14.05) as strong predictors, and *Externalising* remaining inconclusive (BF_incl_ = 0.80). Categorical diagnoses again favoured exclusion (BF_excl_ = 4.21 to 20.00; see Table 2 and Supplementary Sections 5.6, 6.6, and 7.5 for details).

## 4. Discussion

The current study used a naturalistic fMRI ToM paradigm in a sample of psychiatric patients and healthy controls to investigate the transdiagnostic architecture of ToM deficits. Specifically, we aimed to determine (1) whether psychiatric patients as a whole exhibit altered ToM network activation compared to healthy controls, (2) whether these neural responses diverge across five distinct categorical diagnostic clusters (SUD, SZD, AD, ANX, and PD), and (3) whether continuous, transdiagnostic symptom dimensions offer greater predictive utility for ToM-related cortical activation than traditional diagnostic categories. Our main findings were threefold: First, psychiatric patients demonstrated decreased ToM sensitivity in ToM-selective regions relative to healthy controls. Second, we found no evidence for diagnosis-specific activation profiles across the five psychiatric groups. Finally, Bayesian model comparisons revealed that dimensional symptom models provided stronger explanatory power for the observed brain activation patterns than categorical diagnostic groups.

The observed reduction in ToM-specific activation across all examined clinical samples supports the notion that naturalistic paradigms are sensitive and ecologically valid tools that successfully detect specific neural alterations during dynamic social processing while bypassing the cognitive burden of traditional testing (Jääskeläinen & Kosonogov, 2023; Redcay & Moraczewski, 2020; Vanderwal et al., 2019). This finding further aligns with evidence that social-cognitive difficulties often span traditional diagnostic boundaries rather than being restricted to specific categories(Barch, 2020; Cotter et al., 2018a; Dalgleish et al., 2020; De La Higuera-Gonzalez et al., 2023). Rather than reflecting a uniform deficit, individuals across diverse clinical profiles exhibit a less pronounced neural differentiation between ToM and control stimuli than healthy controls. This supports the concept of social brain alterations as a transdiagnostic psychiatric feature (Caspi & Moffitt, 2018).

While behavioural ToM deficits are evident across a wide range of psychiatric diagnoses (Cotter et al., 2018a), similar phenotypic outcomes do not necessarily stem from the same neurobiological causes (Dodell-Feder & Germine, 2018; Henderson et al., 2020). Consequently, prior neuroimaging research relied heavily on case-control paradigms to isolate disorder-specific cortical alterations. However, our lack of diagnosis-specific ToM activation aligns with evidence that purely illness-specific neural signatures are inconsistent (Segal et al., 2023). Instead, widespread transdiagnostic overlaps suggest that traditional categorical nosologies lack the phenotypic resolution to uncover the pathophysiology of social cognition (Segal et al., 2023; Tiego et al., 2023).

To address this, clinical neuroscience is shifting toward dimensional models like the p-factor or the HiTOP (Hierarchical Taxonomy of Psychopathology) framework. These approaches yield robust neuroimaging findings centred on the default mode network (DMN), which heavily overlaps with the ToM network (Porcelli et al., 2019; Schurz et al., 2020). For instance, transdiagnostic dimensions map directly onto DMN hyperconnectivity (Elliott et al., 2018) and share a universal loss of segregation between the DMN and executive networks (Xia et al., 2018). Crucially, this explanatory power extends to clinical outcomes, as dimensional psychopathology significantly outperforms traditional ICD-10 categories in explaining everyday functional disability (Martin et al., 2021; Nagel et al., 2026).

Our preliminary correlation analyses similarly suggest that increased overall psychopathology (across patients and controls) is associated with decreased ToM sensitivity within the examined ToM network. This informed our final analyses, in which we directly compared the predictive utility of dimensional psychopathology and categorical diagnosis clusters for ToM-specific cortical activation. Our Bayesian ANCOVAs robustly identified dimensional measures as better predictors of cortical activation in the examined regions.

This dimensional relationship revealed distinct regional heterogeneity (Park et al., 2022). Activation in the VMPFC showed robust support for Total Problems, an index resembling general symptom burden. This is in line with previous research showing that elevated general psychopathology and emotion dysregulation have been linked to altered social-cognitive processing, including reduced mentalising accuracy, aberrant neural efficiency during Theory of Mind tasks, and disrupted frontoparietal connectivity (Blain et al., 2025; Cotter et al., 2018a; D’Adda et al., 2026; Sharp et al., 2011). In contrast, models incorporating the ASR Internalising scale received strong support in bilateral TPJ, whereas externalising models yielded only anecdotal evidence. This specific association converges with literature linking temporal ToM responses to internalising features such as rumination and anxiety, which are known to suppress TPJ-related mentalising and alter its network connectivity (Cui et al., 2017; Knight et al., 2019). Thus, while medial prefrontal regions may track general psychological burden, temporoparietal nodes appear specifically vulnerable to internalising features.

However, several limitations warrant consideration. First, our inpatient sample reflected natural admission patterns, resulting in an uneven distribution across diagnostic clusters (e.g., a higher representation of ADs relative to PDs or SUDs). While Bayesian analyses enabled us to quantify evidence against group differences despite these variations, this sample imbalance, combined with classifying patients into broad diagnostic clusters rather than finer ICD-10 subcategories, may have limited our sensitivity to detect subtle, disorder-specific effects. Finer diagnostic granularity was not feasible under pragmatic clinical constraints. Second, we did not systematically record medication status or specific comorbidities. Given that pharmacological treatment can modulate regional BOLD signals, future studies with detailed medication protocols are needed to disentangle treatment effects from intrinsic alterations in mentalising.

In conclusion, these findings reinforce that social-cognitive neural networks do not track categorical diagnostic boundaries, but are systematically modulated by continuous dimensions like overall psychopathology (the p-factor) and internalising symptoms (Caspi & Moffitt, 2018; Elliott et al., 2018; Konrad et al., 2024; Xia et al., 2018). Clinically, this supports a shift toward transdiagnostic interventions that target shared symptom profiles rather than specific diagnoses. For example, the Unified Protocol has proven highly efficacious in mitigating core dysregulation across diverse diagnostic groups (Barlow et al., 2017; Caspi & Moffitt, 2018; Kotov et al., 2022), while targeted cognitive-emotional remediation not only improves mentalising deficits but simultaneously alleviates broader psychiatric symptoms (Cotter et al., 2018b; Iacoviello & Charney, 2015). Finally, the strong predictive utility of ASR scores highlights that cost-effective self-report measures can accurately capture this transdiagnostic variance. By measuring individual variance that traditional diagnoses overlook, such dimensional frameworks offer an accessible tool to bridge the gap between complex neurobiological research and real-world clinical practice(Achenbach & Rescorla, 2003; De Vries et al., 2020; Nagel et al., 2026). Further research is needed to determine whether behavioural improvements parallel a normalisation of ToM network activation, and whether adopting such dimensional frameworks can ultimately bridge the translational gap between neuroimaging research and routine clinical care.

## 5. Acknowledgments

We would like to thank Mario Wallner and Franziska Sahm for their help with data acquisition.

## 6. Author Contributions

**P.M.N.** conducted the formal analyses, generated the visualisations, drafted the original manuscript, and helped with data collection. **S.H.** generated the visualisations, helped with the formal analyses, and provided critical revisions to the manuscript. **S.S.-Y.** scripted the fMRI data preprocessing. **D.F.** managed data handling and processing. **K.W.** contributed to manuscript revision. **W.A.** conceptualised the initial study design and facilitated clinical data collection. **M.K.** and **L.K.** conceptualised the initial study design and provided overall supervision. **L.K.** conducted the data collection and formal analyses, refined the theoretical framework and revised the manuscript. All authors reviewed the manuscript.

## 7. Data Availability

The dataset analysed during the current study is available from the corresponding author upon request.

