## Supplementary_material for "Dimensional and Categorical Predictors of Theory of Mind Network Activation: A transdiagnostic fMRI Study in Psychiatric Inpatients"

 

1. Frequency Tables for Educational Level        5

2. Subtext fMRI Methods        11

3. Patient vs Control ANCOVAs        12

3.1 ANCOVA VMPFC        12

3.1.1 Post Hoc Tests        13

3.2 ANCOVA MMPFC        14

3.2.1 Post Hoc Tests        15

3.3 ANCOVA DMPFC        18

3.3.1 Post Hoc Tests        19

3.4 ANCOVA PC        20

3.4.1 Post Hoc Tests        21

3.5 ANCOVA LTPJ        25

3.5.1 Post Hoc Tests        26

3.6 ANCOVA RTPJ        28

3.6.1 Post Hoc Tests        29

4. Between Patient Cluster ANCOVAs        30

4.1 ANCOVA VMPFC        30

4.2 ANCOVA MMPFC        31

4.3 ANCOVA DMPFC        32

4.4 ANCOVA PC        33

4.5 ANCOVA LTPJ        34

4.6 ANCOVA RTPJ        35

5. Bayesian ANCOVAs        36

5.1 Bayesian ANCOVA Internalising VMPFC        36

5.1.1 Analysis of Effects - VMPFC        37

5.1.2 Model Averaged Posterior Summary        38

5.2 Bayesian ANCOVA Internalising MMPFC        42

5.2.1 Analysis of Effects - MMPFC        43

5.2.2 Model Averaged Posterior Summary        44

5.3 Bayesian ANCOVA Internalising DMPFC        48

5.3.1 Analysis of Effects - DMPFC        49

5.3.2 Model Averaged Posterior Summary        50

5.4 Bayesian ANCOVA Internalising PC        54

5.4.1 Analysis of Effects - PC        55

5.4.2 Model Averaged Posterior Summary        56

5.5 Bayesian ANCOVA Internalising LTPJ        60

5.5.1 Analysis of Effects - LTPJ        61

5.5.2 Model Averaged Posterior Summary        62

5.6 Bayesian ANCOVA Internalising RTPJ        66

5.6.1 Analysis of Effects - RTPJ        67

5.6.2 Model Averaged Posterior Summary        68

6. Bayesian ANCOVAs with Externalising as predictor        72

6.1 Bayesian ANCOVA Externalising VMPFC        72

6.1.1 Analysis of Effects - VMPFC        73

6.1.2 Model Averaged Posterior Summary        74

6.2 Bayesian ANCOVA Externalising MMPFC        78

6.2.1 Analysis of Effects - MMPFC        79

6.2.2 Model Averaged Posterior Summary        80

6.3 Bayesian ANCOVA Externalising DMPFC        84

6.3.1 Analysis of Effects - DMPFC        85

6.3.2 Model Averaged Posterior Summary        86

6.4 Bayesian ANCOVA Externalising PC        90

6.4.1 Analysis of Effects - PC        91

6.4.2 Model Averaged Posterior Summary        92

6.5 Bayesian ANCOVA EXT LTPJ        96

6.5.1 Analysis of Effects - LTPJ        97

6.5.2 Model Averaged Posterior Summary        98

6.6 Bayesian ANCOVA Externalising RTPJ        102

6.6.1 Analysis of Effects - RTPJ        103

6.6.2 Model Averaged Posterior Summary        103

7. Bayesian ANCOVAs Total Score        107

7.1 Bayesian ANCOVA Total VMPFC        107

7.1.1 Analysis of Effects - VMPFC        108

7.1.2 Model Averaged Posterior Summary        109

7.2 Bayesian ANCOVA TOTAL MMPFC        113

7.2.1 Analysis of Effects - MMPFC        114

7.2.2 Model Averaged Posterior Summary        115

7.3  Bayesian ANCOVA TOTAL DMPFC        119

7.3.1 Analysis of Effects - DMPFC        120

7.3.2 Model Averaged Posterior Summary        121

7.4 Bayesian ANCOVA TOTAL PC        125

7.4.1 Analysis of Effects - PC        126

7.4.2 Model Averaged Posterior Summary        127

7.5  Bayesian ANCOVA TOTAL RTPJ        131

7.5.1 Analysis of Effects - RTPJ        132

7.5.2 Model Averaged Posterior Summary        133

7.6 Bayesian ANCOVA TOTAL LTPJ        137

7.6.1 Analysis of Effects - LTPJ        138

7.6.2 Model Averaged Posterior Summary        139

---

1. # Frequency Tables for Educational Level

|  |  |  |  |
| --- | --- | --- | --- |
| Frequencies for ISCED-Classification | | | |
| Group | ISCED-Classification | Frequency | Percent |
| Controls | 1 | 0 | 0.0 |
|  | 2 | 4 | 4.4 |
|  | 3 | 53 | 58.2 |
|  | 5 | 0 | 0.0 |
|  | 6 | 33 | 36.3 |
|  | 7 | 0 | 0.0 |
|  | 8 | 1 | 1.1 |
|  | Missing | 0 | 0.0 |
|  | Total | 91 | 100.0 |
| SUD | 1 | 0 | 0.0 |
|  | 2 | 3 | 12.5 |
|  | 3 | 14 | 58.3 |
|  | 5 | 1 | 4.2 |
|  | 6 | 6 | 25.0 |
|  | 7 | 0 | 0.0 |
|  | 8 | 0 | 0.0 |
|  | Missing | 0 | 0.0 |
|  | Total | 24 | 100.0 |
| SSD | 1 | 1 | 2.1 |
|  | 2 | 12 | 25.5 |
|  | 3 | 23 | 48.9 |
|  | 5 | 0 | 0.0 |
|  | 6 | 10 | 21.3 |
|  | 7 | 1 | 2.1 |
|  | 8 | 0 | 0.0 |
|  | Missing | 0 | 0.0 |
|  | Total | 47 | 100.0 |
| AD | 1 | 2 | 2.0 |
|  | 2 | 15 | 14.7 |
|  | 3 | 53 | 52.0 |
|  | 5 | 3 | 2.9 |
|  | 6 | 24 | 23.5 |
|  | 7 | 4 | 3.9 |
|  | 8 | 1 | 1.0 |
|  | Missing | 0 | 0.0 |
|  | Total | 102 | 100.0 |
| ANX | 1 | 0 | 0.0 |
|  | 2 | 6 | 18.2 |
|  | 3 | 17 | 51.5 |
|  | 5 | 0 | 0.0 |
|  | 6 | 9 | 27.3 |
|  | 7 | 1 | 3.0 |
|  | 8 | 0 | 0.0 |
|  | Missing | 0 | 0.0 |
|  | Total | 33 | 100.0 |
| PD | 1 | 1 | 5.3 |
|  | 2 | 9 | 47.4 |
|  | 3 | 8 | 42.1 |
|  | 5 | 0 | 0.0 |
|  | 6 | 0 | 0.0 |
|  | 7 | 1 | 5.3 |
|  | 8 | 0 | 0.0 |
|  | Missing | 0 | 0.0 |
|  | Total | 19 | 100.0 |

---

2. # Subtext fMRI Methods

Functional images were realigned, de-spiked (with the AFNI 3ddespike function), unwarped, corrected for geometric distortions using the fieldmap of each participant, and slice-time corrected. The high-resolution structural T1-weighted image of each participant was processed and normalised with the CAT12 toolbox (http://dbm.neuro.uni-jena.de/cat) using default settings; each structural image was segmented into grey matter, white matter and CSF and denoised, then each image was warped into MNI space by registering it to the DARTEL template provided by the CAT12 toolbox via the high-dimensional DARTEL (Ashburner, 2007) registration algorithm. Based on these steps, a skull-stripped version of each image in native space was created.

To normalise functional images into MNI space, the functional images were coregistered to the skull-stripped structural image and the parameters from the DARTEL registration were used to warp the functional images, which were resampled to  3 x 3 x 3 mm voxels and smoothed with a 6 mm FWHM Gaussian kernel.

 Statistical analysis was performed with a GLM two-stage mixed-effects model. In the subject-specific first level model, each condition was modelled by convolving stick functions at its onsets with SPM12’s canonical hemodynamic response function (target trials and start and end messages were modelled as separate events of no interest, the model also included the six motion parameters and six noise regressors, reflecting physiological noise components obtained from FIACH (Tierney et al., 2016) as regressors of no interest). Parameter estimates for each condition were calculated via these first-level general linear models (GLM), using a temporal high-pass filter (cutoff 128 sec) to remove low-frequency drifts and modelling temporal autocorrelation across scans with an AR (1) process (Friston et al., 2002).

---

3. # Patient vs Control ANCOVAs

### 3.1 ANCOVA VMPFC

|  |  |  |  |  |  |
| --- | --- | --- | --- | --- | --- |
| Cases | Sum of Squares | df | Mean Square | F | p |
| Sex | 22.84 | 1 | 22.84 | 2.135 | .145 |
| Group | 49.74 | 1 | 49.74 | 4.648 | .032 |
| Age | 2.350 | 1 | 2.350 | 0.2197 | .640 |
| Sex ✻ Group | 10.66 | 1 | 10.66 | 0.9965 | .319 |
| Residuals | 3274 | 306 | 10.70 |  |  |
| Note.  Type III Sum of Squares | | | | | |

---

#### 3.1.1 Post Hoc Tests

Standard (HSD)

|  |  |  |  |  |  |  |  |
| --- | --- | --- | --- | --- | --- | --- | --- |
| Post Hoc Comparisons - Group | | | | | | | |
|  |  | Mean Difference | SE | df | t | pTukey | pBonf |
| controls | patients | 0.8815 | 0.4089 | 306 | 2.156 | .032\* | .032\* |
| \* p < .05 | | | | | | | |
| Note.  Results are averaged over the levels of: Sex | | | | | | | |

---

### 3.2 ANCOVA MMPFC

|  |  |  |  |  |  |
| --- | --- | --- | --- | --- | --- |
| Cases | Sum of Squares | df | Mean Square | F | p |
| Sex | 19.76 | 1 | 19.76 | 1.657 | .199 |
| Group | 160.9 | 1 | 160.9 | 13.50 | < .001 |
| Age | 25.71 | 1 | 25.71 | 2.156 | .143 |
| Sex ✻ Group | 63.20 | 1 | 63.20 | 5.301 | .022 |
| Residuals | 3660 | 307 | 11.92 |  |  |
| Note.  Type III Sum of Squares | | | | | |

---

#### 3.2.1 Post Hoc Tests

Standard (HSD)

|  |  |  |  |  |  |  |  |
| --- | --- | --- | --- | --- | --- | --- | --- |
| Post Hoc Comparisons - Group | | | | | | | |
|  |  | Mean Difference | SE | df | t | pTukey | pBonf |
| controls | patients | 1.585 | 0.4313 | 307 | 3.674 | < .001\*\*\* | < .001\*\*\* |
| \*\*\* p < .001 | | | | | | | |
| Note.  Results are averaged over the levels of: Sex | | | | | | | |

|  |  |  |  |  |  |  |  |
| --- | --- | --- | --- | --- | --- | --- | --- |
| Post Hoc Comparisons - Sex ✻ Group | | | | | | | |
|  |  | Mean Difference | SE | df | t | pTukey | pBonf |
| f controls | m controls | -1.552 | 0.7282 | 307 | -2.131 | .146 | .203 |
|  | f patients | 0.5902 | 0.6261 | 307 | 0.9426 | .782 | 1.000 |
|  | m patients | 1.027 | 0.6122 | 307 | 1.678 | .337 | .566 |
| m controls | f patients | 2.142 | 0.6101 | 307 | 3.511 | .003\*\* | .003\*\* |
|  | m patients | 2.579 | 0.5942 | 307 | 4.340 | < .001\*\*\* | < .001\*\*\* |
| f patients | m patients | 0.4371 | 0.4664 | 307 | 0.9371 | .785 | 1.000 |
| \*\* p < .01, \*\*\* p < .001 | | | | | | | |
| Note.  P-value adjusted for comparing a family of 4 estimates. | | | | | | | |

---

### 3.3 ANCOVA DMPFC

|  |  |  |  |  |  |
| --- | --- | --- | --- | --- | --- |
| Cases | Sum of Squares | df | Mean Square | F | p |
| Sex | 8.127 | 1 | 8.127 | 0.7584 | .385 |
| Group | 49.29 | 1 | 49.29 | 4.600 | .033 |
| Age | 13.61 | 1 | 13.61 | 1.270 | .261 |
| Sex ✻ Group | 10.83 | 1 | 10.83 | 1.011 | .315 |
| Residuals | 3290 | 307 | 10.72 |  |  |
| Note.  Type III Sum of Squares | | | | | |

---

#### 3.3.1 Post Hoc Tests

Standard (HSD)

|  |  |  |  |  |  |  |  |
| --- | --- | --- | --- | --- | --- | --- | --- |
| Post Hoc Comparisons - Group | | | | | | | |
|  |  | Mean Difference | SE | df | t | pTukey | pBonf |
| controls | patients | 0.8770 | 0.4089 | 307 | 2.145 | .033\* | .033\* |
| \* p < .05 | | | | | | | |
| Note.  Results are averaged over the levels of: Sex | | | | | | | |

---

#### 3.4 ANCOVA PC

|  |  |  |  |  |  |
| --- | --- | --- | --- | --- | --- |
| Cases | Sum of Squares | df | Mean Square | F | p |
| Sex | 99.15 | 1 | 99.15 | 4.785 | .029 |
| Group | 177.0 | 1 | 177.0 | 8.540 | .004 |
| Age | 132.5 | 1 | 132.5 | 6.395 | .012 |
| Sex ✻ Group | 34.51 | 1 | 34.51 | 1.665 | .198 |
| Residuals | 6362 | 307 | 20.72 |  |  |
| Note.  Type III Sum of Squares | | | | | |

---

#### 3.4.1 Post Hoc Tests

Standard (HSD)

|  |  |  |  |  |  |  |  |
| --- | --- | --- | --- | --- | --- | --- | --- |
| Post Hoc Comparisons - Sex | | | | | | | |
|  |  | Mean Difference | SE | df | t | pTukey | pBonf |
| f | m | -1.248 | 0.5707 | 307 | -2.187 | .029\* | .029\* |
| \* p < .05 | | | | | | | |
| Note.  Results are averaged over the levels of: Group | | | | | | | |

|  |  |  |  |  |  |  |  |
| --- | --- | --- | --- | --- | --- | --- | --- |
| Post Hoc Comparisons - Group | | | | | | | |
|  |  | Mean Difference | SE | df | t | pTukey | pBonf |
| controls | patients | 1.662 | 0.5687 | 307 | 2.922 | .004\*\* | .004\*\* |
| \*\* p < .01 | | | | | | | |
| Note.  Results are averaged over the levels of: Sex | | | | | | | |

|  |  |  |  |  |  |  |  |
| --- | --- | --- | --- | --- | --- | --- | --- |
| Post Hoc Comparisons - Sex ✻ Group | | | | | | | |
|  |  | Mean Difference | SE | df | t | pTukey | pBonf |
| f controls | m controls | -1.983 | 0.9601 | 307 | -2.066 | .167 | .238 |
|  | f patients | 0.9270 | 0.8255 | 307 | 1.123 | .676 | 1.000 |
|  | m patients | 0.4134 | 0.8071 | 307 | 0.5122 | .956 | 1.000 |
| m controls | f patients | 2.910 | 0.8043 | 307 | 3.618 | .002\*\* | .002\*\* |
|  | m controls | 2.397 | 0.7834 | 307 | 3.059 | .013\* | .014\* |
| f controls | m controls | -0.5136 | 0.6149 | 307 | -0.8352 | .838 | 1.000 |
| \* p < .05, \*\* p < .01 | | | | | | | |
| Note.  P-value adjusted for comparing a family of 4 estimates. | | | | | | | |

---

### 3.5 ANCOVA LTPJ

|  |  |  |  |  |  |
| --- | --- | --- | --- | --- | --- |
| Cases | Sum of Squares | df | Mean Square | F | p |
| Sex | 74.29 | 1 | 74.29 | 4.177 | .042 |
| Group | 108.5 | 1 | 108.5 | 6.103 | .014 |
| Age | 8.621 | 1 | 8.621 | 0.4848 | .487 |
| Sex ✻ Group | 11.67 | 1 | 11.67 | 0.6560 | .419 |
| Residuals | 5459 | 307 | 17.78 |  |  |
| Note.  Type III Sum of Squares | | | | | |

---

#### 3.5.1 Post Hoc Tests

Standard (HSD)

|  |  |  |  |  |  |  |  |
| --- | --- | --- | --- | --- | --- | --- | --- |
| Post Hoc Comparisons - Sex | | | | | | | |
|  |  | Mean Difference | SE | df | t | pTukey | pBonf |
| f | m | -1.081 | 0.5287 | 307 | -2.044 | .042\* | .042\* |
| \* p < .05 | | | | | | | |
| Note.  Results are averaged over the levels of: Group | | | | | | | |

|  |  |  |  |  |  |  |  |
| --- | --- | --- | --- | --- | --- | --- | --- |
| Post Hoc Comparisons - Group | | | | | | | |
|  |  | Mean Difference | SE | df | t | pTukey | pBonf |
| controls | patients | 1.301 | 0.5268 | 307 | 2.470 | .014\* | .014\* |
| \* p < .05 | | | | | | | |
| Note.  Results are averaged over the levels of: Sex | | | | | | | |

---

### 3.6 ANCOVA RTPJ

|  |  |  |  |  |  |
| --- | --- | --- | --- | --- | --- |
| Cases | Sum of Squares | df | Mean Square | F | p |
| Sex | 10.75 | 1 | 10.75 | 0.6495 | .421 |
| Group | 75.47 | 1 | 75.47 | 4.558 | .034 |
| Age | 29.87 | 1 | 29.87 | 1.804 | .180 |
| Sex ✻ Group | 10.67 | 1 | 10.67 | 0.6442 | .423 |
| Residuals | 5083 | 307 | 16.56 |  |  |
| Note.  Type III Sum of Squares | | | | | |

---

#### 3.6.1 Post Hoc Tests

Standard (HSD)

|  |  |  |  |  |  |  |  |
| --- | --- | --- | --- | --- | --- | --- | --- |
| Post Hoc Comparisons - Group | | | | | | | |
|  |  | Mean Difference | SE | df | t | pTukey | pBonf |
| controls | patients | 1.085 | 0.5083 | 307 | 2.135 | .034\* | .034\* |
| \* p < .05 | | | | | | | |
| Note.  Results are averaged over the levels of: Sex | | | | | | | |

4. # Between Patient Cluster ANCOVAs

### 4.1 ANCOVA VMPFC

|  |  |  |  |  |  |
| --- | --- | --- | --- | --- | --- |
| Cases | Sum of Squares | df | Mean Square | F | p |
| diagnosis\_cluster | 54.94 | 4 | 13.74 | 1.267 | .284 |
| Sex | 0.8495 | 1 | 0.8495 | 0.07839 | .780 |
| Age | 12.38 | 1 | 12.38 | 1.143 | .286 |
| diagnosis\_cluster ✻ Sex | 0.4633 | 4 | 0.1158 | 0.01069 | 1.000 |
| Residuals | 2265 | 209 | 10.84 |  |  |
| Note.  Type III Sum of Squares | | | | | |

---

### 4.2 ANCOVA MMPFC

|  |  |  |  |  |  |
| --- | --- | --- | --- | --- | --- |
| ANCOVA - MMPFC | | | | | |
| Cases | Sum of Squares | df | Mean Square | F | p |
| diagnosis\_cluster | 49.52 | 4 | 12.38 | 0.9965 | .410 |
| Sex | 12.38 | 1 | 12.38 | 0.9962 | .319 |
| Age | 27.67 | 1 | 27.67 | 2.227 | .137 |
| diagnosis\_cluster ✻ Sex | 21.37 | 4 | 5.344 | 0.4301 | .787 |
| Residuals | 2609 | 210 | 12.42 |  |  |
| Note.  Type III Sum of Squares | | | | | |

---

### 4.3 ANCOVA DMPFC

|  |  |  |  |  |  |
| --- | --- | --- | --- | --- | --- |
| ANCOVA - DMPFC | | | | | |
| Cases | Sum of Squares | df | Mean Square | F | p |
| diagnosis\_cluster | 66.73 | 4 | 16.68 | 1.690 | .153 |
| Sex | 5.727 | 1 | 5.727 | 0.5802 | .447 |
| Age | 50.62 | 1 | 50.62 | 5.129 | .025 |
| diagnosis\_cluster ✻ Sex | 37.99 | 4 | 9.499 | 0.9624 | .429 |
| Residuals | 2073 | 210 | 9.870 |  |  |
| Note.  Type III Sum of Squares | | | | | |

---

### 4.4 ANCOVA PC

|  |  |  |  |  |  |
| --- | --- | --- | --- | --- | --- |
| ANCOVA - PC | | | | | |
| Cases | Sum of Squares | df | Mean Square | F | p |
| diagnosis\_cluster | 51.05 | 4 | 12.76 | 0.6209 | .648 |
| Sex | 3.995 | 1 | 3.995 | 0.1944 | .660 |
| Age | 137.7 | 1 | 137.7 | 6.701 | .010 |
| diagnosis\_cluster ✻ Sex | 79.79 | 4 | 19.95 | 0.9704 | .425 |
| Residuals | 4316 | 210 | 20.55 |  |  |
| Note.  Type III Sum of Squares | | | | | |

---

### 4.5 ANCOVA LTPJ

|  |  |  |  |  |  |
| --- | --- | --- | --- | --- | --- |
| ANCOVA - LTPJ | | | | | |
| Cases | Sum of Squares | df | Mean Square | F | p |
| diagnosis\_cluster | 126.1 | 4 | 31.51 | 1.831 | .124 |
| Sex | 3.357 | 1 | 3.357 | 0.1950 | .659 |
| Age | 27.79 | 1 | 27.79 | 1.614 | .205 |
| diagnosis\_cluster ✻ Sex | 132.2 | 4 | 33.04 | 1.919 | .108 |
| Residuals | 3615 | 210 | 17.21 |  |  |
| Note.  Type III Sum of Squares | | | | | |

---

### 4.6 ANCOVA RTPJ

|  |  |  |  |  |  |
| --- | --- | --- | --- | --- | --- |
| ANCOVA - RTPJ | | | | | |
| Cases | Sum of Squares | df | Mean Square | F | p |
| diagnosis\_cluster | 67.17 | 4 | 16.79 | 0.9675 | .426 |
| Sex | 15.41 | 1 | 15.41 | 0.8876 | .347 |
| Age | 79.63 | 1 | 79.63 | 4.588 | .033 |
| diagnosis\_cluster ✻ Sex | 35.79 | 4 | 8.947 | 0.5155 | .724 |
| Residuals | 3645 | 210 | 17.36 |  |  |
| Note.  Type III Sum of Squares | | | | | |

---

5. # Bayesian ANCOVAs

### 5.1 Bayesian ANCOVA Internalising VMPFC

|  |  |  |  |  |  |
| --- | --- | --- | --- | --- | --- |
| Model Comparison | | | | | |
| Models | P(M) | P(M\|data) | BFM | BF10 | error % |
| ASR\_Internalizing | 0.1667 | 0.8809 | 36.99 | 1.000 |  |
| diagnosis\_cluster + ASR\_Internalizing | 0.1667 | 0.09028 | 0.4962 | 0.1025 | 2.863 |
| diagnosis\_cluster | 0.1667 | 0.01514 | 0.07685 | 0.01718 | 2.826 |
| Null model (incl. Sex, Age) | 0.1667 | 0.008167 | 0.04117 | 0.009270 | 1.791 |
| diagnosis\_cluster + ASR\_Internalizing + diagnosis\_cluster ✻  Sex | 0.1667 | 0.004815 | 0.02419 | 0.005465 | 4.242 |
| diagnosis\_cluster + diagnosis\_cluster ✻  Sex | 0.1667 | 6.593×10-4 | 0.003299 | 7.484×10-4 | 3.847 |
| Note.  All models include Sex, Age. | | | | | |

|  |  |  |  |  |  |
| --- | --- | --- | --- | --- | --- |
| 5.1.1 Analysis of Effects - VMPFC | | | | | |
| Effects | P(incl) | P(excl) | P(incl\|data) | P(excl\|data) | BFincl |
| diagnosis\_cluster | 0.6667 | 0.3333 | 0.1109 | 0.8891 | 0.06236 |
| diagnosis\_cluster ✻  Sex | 0.3333 | 0.6667 | 0.005474 | 0.9945 | 0.01101 |
| ASR\_Internalizing | 0.5000 | 0.5000 | 0.9760 | 0.02396 | 40.73 |

|  |  |  |  |  |  |
| --- | --- | --- | --- | --- | --- |
| 5.1.2 Model Averaged Posterior Summary | | | | | |
|  | | | | 95% Credible Interval | |
| Variable | Level | Mean | SD | Lower | Upper |
| Intercept |  | 1.898 | 0.1976 | 1.502 | 2.288 |
| diagnosis\_cluster | 1 | -0.3163 | 0.5034 | -1.330 | 0.6769 |
|  | 2 | -0.2756 | 0.4315 | -1.152 | 0.5845 |
|  | 3 | -0.2851 | 0.3398 | -0.9756 | 0.3887 |
|  | 4 | 0.7566 | 0.4793 | -0.1622 | 1.757 |
|  | 6 | -0.3128 | 0.5775 | -1.512 | 0.8050 |
|  | 9 | 0.4331 | 0.4154 | -0.3715 | 1.287 |
| ASR\_Internalizing |  | -0.04657 | 0.01370 | -0.07552 | -0.02120 |
| Sex | f | -0.03363 | 0.1938 | -0.4200 | 0.3516 |
|  | m | 0.03363 | 0.1938 | -0.3548 | 0.4172 |
| Age |  | -0.01420 | 0.01387 | -0.04653 | 0.01258 |
| diagnosis\_cluster ✻  Sex | 1 & f | -0.001593 | 0.4746 | -0.9583 | 0.9430 |
|  | 1 & m | 0.001593 | 0.4746 | -0.9514 | 0.9498 |
|  | 2 & f | 0.1307 | 0.3872 | -0.6381 | 0.9075 |
|  | 2 & m | -0.1307 | 0.3872 | -0.9180 | 0.6271 |
|  | 3 & f | -0.02460 | 0.3068 | -0.6413 | 0.5785 |
|  | 3 & m | 0.02460 | 0.3068 | -0.5864 | 0.6342 |
|  | 4 & f | 0.08561 | 0.4048 | -0.7200 | 0.9006 |
|  | 4 & m | -0.08561 | 0.4048 | -0.9140 | 0.7146 |
|  | 6 & f | 0.1490 | 0.4602 | -0.7493 | 1.096 |
|  | 6 & m | -0.1490 | 0.4602 | -1.110 | 0.7331 |
|  | 9 & f | -0.3391 | 0.3211 | -1.005 | 0.2902 |
|  | 9 & m | 0.3391 | 0.3211 | -0.2997 | 0.9999 |

---

### 5.2 Bayesian ANCOVA Internalising MMPFC

|  |  |  |  |  |  |
| --- | --- | --- | --- | --- | --- |
| Model Comparison | | | | | |
| Models | P(M) | P(M\|data) | BFM | BF10 | error % |
| ASR\_Internalizing | 0.1667 | 0.4335 | 3.826 | 1.000 |  |
| diagnosis\_cluster | 0.1667 | 0.1932 | 1.198 | 0.4458 | 8.418 |
| diagnosis\_cluster + ASR\_Internalizing | 0.1667 | 0.1582 | 0.9396 | 0.3649 | 7.303 |
| diagnosis\_cluster + ASR\_Internalizing + diagnosis\_cluster ✻  Sex | 0.1667 | 0.1090 | 0.6116 | 0.2514 | 7.267 |
| diagnosis\_cluster + diagnosis\_cluster ✻  Sex | 0.1667 | 0.09161 | 0.5042 | 0.2113 | 7.286 |
| Null model (incl. Sex, Age) | 0.1667 | 0.01448 | 0.07347 | 0.03341 | 5.120 |
| Note.  All models include Sex, Age. | | | | | |

|  |  |  |  |  |  |
| --- | --- | --- | --- | --- | --- |
| 5.2.1 Analysis of Effects - MMPFC | | | | | |
| Effects | P(incl) | P(excl) | P(incl\|data) | P(excl\|data) | BFincl |
| diagnosis\_cluster | 0.6667 | 0.3333 | 0.5520 | 0.4480 | 0.6162 |
| diagnosis\_cluster ✻  Sex | 0.3333 | 0.6667 | 0.2006 | 0.7994 | 0.5019 |
| ASR\_Internalizing | 0.5000 | 0.5000 | 0.7007 | 0.2993 | 2.341 |

|  |  |  |  |  |  |
| --- | --- | --- | --- | --- | --- |
| 5.2.2 Model Averaged Posterior Summary | | | | | |
|  | | | | 95% Credible Interval | |
| Variable | Level | Mean | SD | Lower | Upper |
| Intercept |  | 1.410 | 0.2374 | 0.9234 | 1.867 |
| diagnosis\_cluster | 1 | -0.3284 | 0.5729 | -1.490 | 0.8175 |
|  | 2 | -0.1410 | 0.4688 | -1.083 | 0.7852 |
|  | 3 | -0.4696 | 0.3659 | -1.201 | 0.2546 |
|  | 4 | 0.5185 | 0.4990 | -0.4526 | 1.540 |
|  | 6 | -0.6110 | 0.6358 | -1.928 | 0.5993 |
|  | 9 | 1.032 | 0.4376 | 0.1402 | 1.875 |
| ASR\_Internalizing |  | -0.03874 | 0.01762 | -0.07258 | -0.002824 |
| Sex | f | 0.1146 | 0.2254 | -0.3265 | 0.5705 |
|  | m | -0.1146 | 0.2254 | -0.5745 | 0.3252 |
| Age |  | -0.02373 | 0.01546 | -0.05598 | -0.002379 |
| diagnosis\_cluster ✻  Sex | 1 & f | 0.1529 | 0.5501 | -0.9666 | 1.270 |
|  | 1 & m | -0.1529 | 0.5501 | -1.282 | 0.9521 |
|  | 2 & f | 0.03558 | 0.4320 | -0.8300 | 0.8996 |
|  | 2 & m | -0.03558 | 0.4320 | -0.9081 | 0.8215 |
|  | 3 & f | 0.1409 | 0.3390 | -0.5386 | 0.8177 |
|  | 3 & m | -0.1409 | 0.3390 | -0.8227 | 0.5313 |
|  | 4 & f | -0.07350 | 0.4500 | -0.9814 | 0.8154 |
|  | 4 & m | 0.07350 | 0.4500 | -0.8302 | 0.9718 |
|  | 6 & f | 0.5776 | 0.5580 | -0.4862 | 1.764 |
|  | 6 & m | -0.5776 | 0.5580 | -1.770 | 0.4649 |
|  | 9 & f | -0.8334 | 0.3743 | -1.603 | -0.1226 |
|  | 9 & m | 0.8334 | 0.3743 | 0.1089 | 1.598 |

---

### 5.3 Bayesian ANCOVA Internalising DMPFC

|  |  |  |  |  |  |
| --- | --- | --- | --- | --- | --- |
| Model Comparison | | | | | |
| Models | P(M) | P(M\|data) | BFM | BF10 | error % |
| ASR\_Internalizing | 0.1667 | 0.4215 | 3.643 | 1.000 |  |
| diagnosis\_cluster | 0.1667 | 0.2095 | 1.325 | 0.4970 | 5.439 |
| Null model (incl. Sex, Age) | 0.1667 | 0.1905 | 1.176 | 0.4519 | 3.597 |
| diagnosis\_cluster + ASR\_Internalizing | 0.1667 | 0.1527 | 0.9011 | 0.3623 | 4.910 |
| diagnosis\_cluster + diagnosis\_cluster ✻  Sex | 0.1667 | 0.01420 | 0.07201 | 0.03368 | 4.897 |
| diagnosis\_cluster + ASR\_Internalizing + diagnosis\_cluster ✻  Sex | 0.1667 | 0.01165 | 0.05892 | 0.02763 | 4.667 |
| Note.  All models include Sex, Age. | | | | | |

|  |  |  |  |  |  |
| --- | --- | --- | --- | --- | --- |
| 5.3.1 Analysis of Effects - DMPFC | | | | | |
| Effects | P(incl) | P(excl) | P(incl\|data) | P(excl\|data) | BFincl |
| diagnosis\_cluster | 0.6667 | 0.3333 | 0.3880 | 0.6120 | 0.3170 |
| diagnosis\_cluster ✻  Sex | 0.3333 | 0.6667 | 0.02584 | 0.9742 | 0.05306 |
| ASR\_Internalizing | 0.5000 | 0.5000 | 0.5859 | 0.4141 | 1.415 |

|  |  |  |  |  |  |
| --- | --- | --- | --- | --- | --- |
| 5.3.2 Model Averaged Posterior Summary | | | | | |
|  | | | | 95% Credible Interval | |
| Variable | Level | Mean | SD | Lower | Upper |
| Intercept |  | 1.980 | 0.2138 | 1.540 | 2.395 |
| diagnosis\_cluster | 1 | -0.08728 | 0.5081 | -1.111 | 0.9201 |
|  | 2 | -0.9018 | 0.4516 | -1.828 | -0.02421 |
|  | 3 | -0.09389 | 0.3401 | -0.7735 | 0.5870 |
|  | 4 | 0.5990 | 0.4703 | -0.3313 | 1.554 |
|  | 6 | -0.1834 | 0.5734 | -1.343 | 0.9559 |
|  | 9 | 0.6674 | 0.3912 | -0.1299 | 1.432 |
| ASR\_Internalizing |  | -0.02749 | 0.01443 | -0.05690 | -2.614×10-4 |
| Sex | f | 0.1132 | 0.1996 | -0.2823 | 0.5116 |
|  | m | -0.1132 | 0.1996 | -0.5132 | 0.2792 |
| Age |  | -0.02104 | 0.01432 | -0.05217 | -0.001461 |
| diagnosis\_cluster ✻  Sex | 1 & f | 0.2139 | 0.5001 | -0.7729 | 1.253 |
|  | 1 & m | -0.2139 | 0.5001 | -1.275 | 0.7575 |
|  | 2 & f | -0.2101 | 0.4043 | -1.042 | 0.5742 |
|  | 2 & m | 0.2101 | 0.4043 | -0.5861 | 1.038 |
|  | 3 & f | -0.1719 | 0.3165 | -0.8204 | 0.4456 |
|  | 3 & m | 0.1719 | 0.3165 | -0.4471 | 0.8196 |
|  | 4 & f | 0.1318 | 0.4132 | -0.7021 | 0.9644 |
|  | 4 & m | -0.1318 | 0.4132 | -0.9710 | 0.6961 |
|  | 6 & f | 0.3805 | 0.4951 | -0.5657 | 1.422 |
|  | 6 & m | -0.3805 | 0.4951 | -1.435 | 0.5599 |
|  | 9 & f | -0.3441 | 0.3303 | -1.030 | 0.2891 |
|  | 9 & m | 0.3441 | 0.3303 | -0.2978 | 1.026 |

---

### 5.4 Bayesian ANCOVA Internalising PC

|  |  |  |  |  |  |
| --- | --- | --- | --- | --- | --- |
| Model Comparison | | | | | |
| Models | P(M) | P(M\|data) | BFM | BF10 | error % |
| ASR\_Internalizing | 0.1667 | 0.4204 | 3.627 | 1.000 |  |
| diagnosis\_cluster | 0.1667 | 0.2976 | 2.119 | 0.7079 | 8.709 |
| diagnosis\_cluster + ASR\_Internalizing | 0.1667 | 0.2000 | 1.250 | 0.4757 | 7.170 |
| diagnosis\_cluster + diagnosis\_cluster ✻  Sex | 0.1667 | 0.02997 | 0.1545 | 0.07129 | 7.013 |
| Null model (incl. Sex, Age) | 0.1667 | 0.02863 | 0.1474 | 0.06811 | 4.812 |
| diagnosis\_cluster + ASR\_Internalizing + diagnosis\_cluster ✻  Sex | 0.1667 | 0.02336 | 0.1196 | 0.05557 | 6.798 |
| Note.  All models include Sex, Age. | | | | | |

|  |  |  |  |  |  |
| --- | --- | --- | --- | --- | --- |
| 5.4.1 Analysis of Effects - PC | | | | | |
| Effects | P(incl) | P(excl) | P(incl\|data) | P(excl\|data) | BFincl |
| diagnosis\_cluster | 0.6667 | 0.3333 | 0.5509 | 0.4491 | 0.6134 |
| diagnosis\_cluster ✻  Sex | 0.3333 | 0.6667 | 0.05334 | 0.9467 | 0.1127 |
| ASR\_Internalizing | 0.5000 | 0.5000 | 0.6438 | 0.3562 | 1.807 |

|  |  |  |  |  |  |
| --- | --- | --- | --- | --- | --- |
| 5.4.2 Model Averaged Posterior Summary | | | | | |
|  | | | | 95% Credible Interval | |
| Variable | Level | Mean | SD | Lower | Upper |
| Intercept |  | 5.217 | 0.3311 | 4.529 | 5.830 |
| diagnosis\_cluster | 1 | -0.9157 | 0.7208 | -2.391 | 0.4890 |
|  | 2 | -0.2390 | 0.5949 | -1.413 | 0.9616 |
|  | 3 | -0.06376 | 0.4577 | -0.9870 | 0.8431 |
|  | 4 | 0.5533 | 0.6337 | -0.6946 | 1.819 |
|  | 6 | -0.8306 | 0.7924 | -2.465 | 0.6953 |
|  | 9 | 1.496 | 0.5512 | 0.3739 | 2.565 |
| ASR\_Internalizing |  | -0.04600 | 0.02195 | -0.09026 | -0.001145 |
| Sex | f | -0.2763 | 0.2761 | -0.8291 | 0.2719 |
|  | m | 0.2763 | 0.2761 | -0.2757 | 0.8254 |
| Age |  | -0.04910 | 0.01951 | -0.09464 | -0.01096 |
| diagnosis\_cluster ✻  Sex | 1 & f | 0.4397 | 0.6903 | -0.8895 | 1.895 |
|  | 1 & m | -0.4397 | 0.6903 | -1.913 | 0.8682 |
|  | 2 & f | -0.3905 | 0.5439 | -1.532 | 0.6500 |
|  | 2 & m | 0.3905 | 0.5439 | -0.6660 | 1.515 |
|  | 3 & f | 0.06642 | 0.4205 | -0.7757 | 0.9011 |
|  | 3 & m | -0.06642 | 0.4205 | -0.9140 | 0.7711 |
|  | 4 & f | 0.2198 | 0.5542 | -0.8842 | 1.355 |
|  | 4 & m | -0.2198 | 0.5542 | -1.360 | 0.8690 |
|  | 6 & f | 0.2786 | 0.6552 | -1.028 | 1.624 |
|  | 6 & m | -0.2786 | 0.6552 | -1.639 | 1.022 |
|  | 9 & f | -0.6139 | 0.4509 | -1.547 | 0.2542 |
|  | 9 & m | 0.6139 | 0.4509 | -0.2608 | 1.535 |

---

### 5.5 Bayesian ANCOVA Internalising LTPJ

|  |  |  |  |  |  |
| --- | --- | --- | --- | --- | --- |
| Model Comparison | | | | | |
| Models | P(M) | P(M\|data) | BFM | BF10 | error % |
| ASR\_Internalizing | 0.1667 | 0.7067 | 12.05 | 1.000 |  |
| diagnosis\_cluster + ASR\_Internalizing | 0.1667 | 0.1703 | 1.026 | 0.2410 | 70.79 |
| diagnosis\_cluster | 0.1667 | 0.04814 | 0.2529 | 0.06811 | 70.84 |
| diagnosis\_cluster + ASR\_Internalizing + diagnosis\_cluster ✻  Sex | 0.1667 | 0.03854 | 0.2004 | 0.05453 | 70.83 |
| Null model (incl. Sex, Age) | 0.1667 | 0.02873 | 0.1479 | 0.04065 | 50.06 |
| diagnosis\_cluster + diagnosis\_cluster ✻  Sex | 0.1667 | 0.007531 | 0.03794 | 0.01066 | 70.80 |
| Note.  All models include Sex, Age. | | | | | |

|  |  |  |  |  |  |
| --- | --- | --- | --- | --- | --- |
| 5.5.1 Analysis of Effects - LTPJ | | | | | |
| Effects | P(incl) | P(excl) | P(incl\|data) | P(excl\|data) | BFincl |
| diagnosis\_cluster | 0.6667 | 0.3333 | 0.2645 | 0.7355 | 0.1798 |
| diagnosis\_cluster ✻  Sex | 0.3333 | 0.6667 | 0.04607 | 0.9539 | 0.09659 |
| ASR\_Internalizing | 0.5000 | 0.5000 | 0.9156 | 0.08440 | 10.85 |

|  |  |  |  |  |  |
| --- | --- | --- | --- | --- | --- |
| 5.5.2 Model Averaged Posterior Summary | | | | | |
|  | | | | 95% Credible Interval | |
| Variable | Level | Mean | SD | Lower | Upper |
| Intercept |  | 3.960 | 0.2573 | 3.439 | 4.466 |
| diagnosis\_cluster | 1 | 0.2799 | 0.6604 | -1.021 | 1.639 |
|  | 2 | -1.012 | 0.5654 | -2.175 | 0.08309 |
|  | 3 | -0.3109 | 0.4325 | -1.189 | 0.5338 |
|  | 4 | 0.7304 | 0.5793 | -0.3984 | 1.903 |
|  | 6 | -0.3717 | 0.7521 | -1.942 | 1.103 |
|  | 9 | 0.6842 | 0.5281 | -0.3570 | 1.739 |
| ASR\_Internalizing |  | -0.05373 | 0.01817 | -0.09178 | -0.02166 |
| Sex | f | -0.1279 | 0.2537 | -0.6327 | 0.3781 |
|  | m | 0.1279 | 0.2537 | -0.3823 | 0.6254 |
| Age |  | -0.01714 | 0.01771 | -0.06357 | 0.007520 |
| diagnosis\_cluster ✻  Sex | 1 & f | 0.3394 | 0.6716 | -0.9562 | 1.733 |
|  | 1 & m | -0.3394 | 0.6716 | -1.749 | 0.9506 |
|  | 2 & f | -0.2645 | 0.5236 | -1.347 | 0.7469 |
|  | 2 & m | 0.2645 | 0.5236 | -0.7531 | 1.338 |
|  | 3 & f | -0.2253 | 0.4095 | -1.077 | 0.5749 |
|  | 3 & m | 0.2253 | 0.4095 | -0.5806 | 1.060 |
|  | 4 & f | -0.2859 | 0.5522 | -1.442 | 0.7779 |
|  | 4 & m | 0.2859 | 0.5522 | -0.7830 | 1.437 |
|  | 6 & f | 1.036 | 0.7181 | -0.2733 | 2.584 |
|  | 6 & m | -1.036 | 0.7181 | -2.597 | 0.2462 |
|  | 9 & f | -0.5999 | 0.4310 | -1.486 | 0.2291 |
|  | 9 & m | 0.5999 | 0.4310 | -0.2335 | 1.480 |

---

### 5.6 Bayesian ANCOVA Internalising RTPJ

|  |  |  |  |  |  |
| --- | --- | --- | --- | --- | --- |
| Model Comparison | | | | | |
| Models | P(M) | P(M\|data) | BFM | BF10 | error % |
| ASR\_Internalizing | 0.1667 | 0.8429 | 26.82 | 1.000 |  |
| diagnosis\_cluster + ASR\_Internalizing | 0.1667 | 0.08790 | 0.4818 | 0.1043 | 3.629 |
| Null model (incl. Sex, Age) | 0.1667 | 0.03722 | 0.1933 | 0.04416 | 1.642 |
| diagnosis\_cluster | 0.1667 | 0.02712 | 0.1394 | 0.03218 | 3.445 |
| diagnosis\_cluster + ASR\_Internalizing + diagnosis\_cluster ✻  Sex | 0.1667 | 0.003666 | 0.01840 | 0.004349 | 3.016 |
| diagnosis\_cluster + diagnosis\_cluster ✻  Sex | 0.1667 | 0.001238 | 0.006196 | 0.001468 | 5.907 |
| Note.  All models include Sex, Age. | | | | | |

|  |  |  |  |  |  |
| --- | --- | --- | --- | --- | --- |
| 5.6.1 Analysis of Effects - RTPJ | | | | | |
| Effects | P(incl) | P(excl) | P(incl\|data) | P(excl\|data) | BFincl |
| diagnosis\_cluster | 0.6667 | 0.3333 | 0.1199 | 0.8801 | 0.06813 |
| diagnosis\_cluster ✻  Sex | 0.3333 | 0.6667 | 0.004903 | 0.9951 | 0.009855 |
| ASR\_Internalizing | 0.5000 | 0.5000 | 0.9344 | 0.06558 | 14.25 |

|  |  |  |  |  |  |
| --- | --- | --- | --- | --- | --- |
| 5.6.2 Model Averaged Posterior Summary | | | | | |
|  | | | | 95% Credible Interval | |
| Variable | Level | Mean | SD | Lower | Upper |
| Intercept |  | 3.685 | 0.2418 | 3.191 | 4.160 |
| diagnosis\_cluster | 1 | 0.1478 | 0.6007 | -1.046 | 1.367 |
|  | 2 | -0.7013 | 0.5334 | -1.794 | 0.3443 |
|  | 3 | 0.07887 | 0.4093 | -0.7450 | 0.8904 |
|  | 4 | 0.7384 | 0.5612 | -0.3491 | 1.890 |
|  | 6 | -0.7820 | 0.7258 | -2.299 | 0.5862 |
|  | 9 | 0.5182 | 0.5076 | -0.4764 | 1.532 |
| ASR\_Internalizing |  | -0.04877 | 0.01653 | -0.08680 | -0.01696 |
| Sex | f | -0.003863 | 0.2358 | -0.4763 | 0.4636 |
|  | m | 0.003863 | 0.2358 | -0.4656 | 0.4689 |
| Age |  | -0.02908 | 0.01686 | -0.07218 | -0.004763 |
| diagnosis\_cluster ✻  Sex | 1 & f | 0.1013 | 0.5872 | -1.057 | 1.310 |
|  | 1 & m | -0.1013 | 0.5872 | -1.324 | 1.054 |
|  | 2 & f | -0.1927 | 0.4653 | -1.152 | 0.7221 |
|  | 2 & m | 0.1927 | 0.4653 | -0.7289 | 1.140 |
|  | 3 & f | 0.1585 | 0.3677 | -0.5706 | 0.8906 |
|  | 3 & m | -0.1585 | 0.3677 | -0.9018 | 0.5631 |
|  | 4 & f | -0.3066 | 0.4883 | -1.338 | 0.6433 |
|  | 4 & m | 0.3066 | 0.4883 | -0.6540 | 1.319 |
|  | 6 & f | 0.1672 | 0.5580 | -0.9504 | 1.325 |
|  | 6 & m | -0.1672 | 0.5580 | -1.338 | 0.9330 |
|  | 9 & f | 0.07232 | 0.3767 | -0.6775 | 0.8248 |
|  | 9 & m | -0.07232 | 0.3767 | -0.8330 | 0.6693 |

---

6. # Bayesian ANCOVAs with Externalising as predictor

### 6.1 Bayesian ANCOVA Externalising VMPFC

|  |  |  |  |  |  |
| --- | --- | --- | --- | --- | --- |
| Model Comparison | | | | | |
| Models | P(M) | P(M\|data) | BFM | BF10 | error % |
| ASR\_Externalizing | 0.1667 | 0.8580 | 30.21 | 1.000 |  |
| diagnosis\_cluster + ASR\_Externalizing | 0.1667 | 0.1174 | 0.6649 | 0.1368 | 4.063 |
| diagnosis\_cluster | 0.1667 | 0.01192 | 0.06031 | 0.01389 | 6.841 |
| diagnosis\_cluster + ASR\_Externalizing + diagnosis\_cluster ✻  Sex | 0.1667 | 0.006467 | 0.03254 | 0.007537 | 6.662 |
| Null model (incl. Sex, Age) | 0.1667 | 0.005763 | 0.02898 | 0.006717 | 1.882 |
| diagnosis\_cluster + diagnosis\_cluster ✻  Sex | 0.1667 | 4.911×10-4 | 0.002457 | 5.724×10-4 | 3.606 |
| Note.  All models include Sex, Age. | | | | | |

|  |  |  |  |  |  |
| --- | --- | --- | --- | --- | --- |
| 6.1.1 Analysis of Effects - VMPFC | | | | | |
| Effects | P(incl) | P(excl) | P(incl\|data) | P(excl\|data) | BFincl |
| diagnosis\_cluster | 0.6667 | 0.3333 | 0.1362 | 0.8638 | 0.07887 |
| diagnosis\_cluster ✻  Sex | 0.3333 | 0.6667 | 0.006958 | 0.9930 | 0.01401 |
| ASR\_Externalizing | 0.5000 | 0.5000 | 0.9818 | 0.01817 | 54.03 |

|  |  |  |  |  |  |
| --- | --- | --- | --- | --- | --- |
| 6.1.2 Model Averaged Posterior Summary | | | | | |
|  | | | | 95% Credible Interval | |
| Variable | Level | Mean | SD | Lower | Upper |
| Intercept |  | 1.889 | 0.1970 | 1.490 | 2.280 |
| diagnosis\_cluster | 1 | -0.1432 | 0.4982 | -1.145 | 0.8343 |
|  | 2 | -0.2335 | 0.4181 | -1.092 | 0.5990 |
|  | 3 | -0.4227 | 0.3281 | -1.086 | 0.2272 |
|  | 4 | 0.6134 | 0.4608 | -0.2828 | 1.543 |
|  | 6 | -0.3235 | 0.5542 | -1.471 | 0.7440 |
|  | 9 | 0.5095 | 0.3709 | -0.2241 | 1.260 |
| ASR\_Externalizing |  | -0.07239 | 0.02139 | -0.1166 | -0.03164 |
| Sex | f | -0.1804 | 0.1898 | -0.5605 | 0.1905 |
|  | m | 0.1804 | 0.1898 | -0.1983 | 0.5563 |
| Age |  | -0.01372 | 0.01388 | -0.05213 | 0.008311 |
| diagnosis\_cluster ✻  Sex | 1 & f | -0.04053 | 0.4798 | -1.026 | 0.9083 |
|  | 1 & m | 0.04053 | 0.4798 | -0.9141 | 1.017 |
|  | 2 & f | 0.2179 | 0.3905 | -0.5540 | 1.016 |
|  | 2 & m | -0.2179 | 0.3905 | -1.028 | 0.5416 |
|  | 3 & f | -0.02148 | 0.3013 | -0.6331 | 0.5729 |
|  | 3 & m | 0.02148 | 0.3013 | -0.5788 | 0.6293 |
|  | 4 & f | 0.1146 | 0.4084 | -0.7099 | 0.9340 |
|  | 4 & m | -0.1146 | 0.4084 | -0.9350 | 0.7005 |
|  | 6 & f | 0.04299 | 0.4589 | -0.8874 | 0.9657 |
|  | 6 & m | -0.04299 | 0.4589 | -0.9776 | 0.8733 |
|  | 9 & f | -0.3134 | 0.3165 | -0.9726 | 0.2892 |
|  | 9 & m | 0.3134 | 0.3165 | -0.3008 | 0.9670 |

---

### 6.2 Bayesian ANCOVA Externalising MMPFC

|  |  |  |  |  |  |
| --- | --- | --- | --- | --- | --- |
| Model Comparison | | | | | |
| Models | P(M) | P(M\|data) | BFM | BF10 | error % |
| ASR\_Externalizing | 0.1667 | 0.4213 | 3.640 | 1.000 |  |
| diagnosis\_cluster + ASR\_Externalizing | 0.1667 | 0.2534 | 1.697 | 0.6014 | 3.103 |
| diagnosis\_cluster + ASR\_Externalizing + diagnosis\_cluster ✻  Sex | 0.1667 | 0.1559 | 0.9236 | 0.3701 | 6.519 |
| diagnosis\_cluster | 0.1667 | 0.1053 | 0.5886 | 0.2500 | 3.703 |
| diagnosis\_cluster + diagnosis\_cluster ✻  Sex | 0.1667 | 0.05592 | 0.2962 | 0.1327 | 5.800 |
| Null model (incl. Sex, Age) | 0.1667 | 0.008166 | 0.04117 | 0.01938 | 2.641 |
| Note.  All models include Sex, Age. | | | | | |

|  |  |  |  |  |  |
| --- | --- | --- | --- | --- | --- |
| 6.2.1 Analysis of Effects - MMPFC | | | | | |
| Effects | P(incl) | P(excl) | P(incl\|data) | P(excl\|data) | BFincl |
| diagnosis\_cluster | 0.6667 | 0.3333 | 0.5705 | 0.4295 | 0.6642 |
| diagnosis\_cluster ✻  Sex | 0.3333 | 0.6667 | 0.2118 | 0.7882 | 0.5376 |
| ASR\_Externalizing | 0.5000 | 0.5000 | 0.8306 | 0.1694 | 4.903 |

|  |  |  |  |  |  |
| --- | --- | --- | --- | --- | --- |
| 6.2.2 Model Averaged Posterior Summary | | | | | |
|  | | | | 95% Credible Interval | |
| Variable | Level | Mean | SD | Lower | Upper |
| Intercept |  | 1.415 | 0.2329 | 0.9340 | 1.868 |
| diagnosis\_cluster | 1 | -0.2332 | 0.5689 | -1.388 | 0.8892 |
|  | 2 | -0.1541 | 0.4635 | -1.086 | 0.7686 |
|  | 3 | -0.5209 | 0.3555 | -1.236 | 0.1778 |
|  | 4 | 0.4475 | 0.4915 | -0.5195 | 1.448 |
|  | 6 | -0.5063 | 0.6353 | -1.841 | 0.6980 |
|  | 9 | 0.9670 | 0.4180 | 0.1487 | 1.816 |
| ASR\_Externalizing |  | -0.06353 | 0.02526 | -0.1135 | -0.01416 |
| Sex | f | 0.02284 | 0.2201 | -0.4031 | 0.4744 |
|  | m | -0.02284 | 0.2201 | -0.4807 | 0.3986 |
| Age |  | -0.02468 | 0.01514 | -0.06371 | -0.003147 |
| diagnosis\_cluster ✻  Sex | 1 & f | 0.1025 | 0.5470 | -1.002 | 1.202 |
|  | 1 & m | -0.1025 | 0.5470 | -1.212 | 0.9874 |
|  | 2 & f | 0.1049 | 0.4301 | -0.7619 | 0.9597 |
|  | 2 & m | -0.1049 | 0.4301 | -0.9727 | 0.7489 |
|  | 3 & f | 0.1454 | 0.3348 | -0.5173 | 0.8164 |
|  | 3 & m | -0.1454 | 0.3348 | -0.8231 | 0.5118 |
|  | 4 & f | -0.04296 | 0.4507 | -0.9723 | 0.8431 |
|  | 4 & m | 0.04296 | 0.4507 | -0.8611 | 0.9643 |
|  | 6 & f | 0.4958 | 0.5435 | -0.5427 | 1.636 |
|  | 6 & m | -0.4958 | 0.5435 | -1.652 | 0.5318 |
|  | 9 & f | -0.8056 | 0.3704 | -1.574 | -0.1039 |
|  | 9 & m | 0.8056 | 0.3704 | 0.08941 | 1.564 |

---

### 6.3 Bayesian ANCOVA Externalising DMPFC

|  |  |  |  |  |  |
| --- | --- | --- | --- | --- | --- |
| Model Comparison | | | | | |
| Models | P(M) | P(M\|data) | BFM | BF10 | error % |
| ASR\_Externalizing | 0.1667 | 0.5626 | 6.431 | 1.000 |  |
| diagnosis\_cluster + ASR\_Externalizing | 0.1667 | 0.1787 | 1.088 | 0.3175 | 8.016 |
| Null model (incl. Sex, Age) | 0.1667 | 0.1236 | 0.7049 | 0.2196 | 5.206 |
| diagnosis\_cluster | 0.1667 | 0.1156 | 0.6536 | 0.2055 | 7.394 |
| diagnosis\_cluster + ASR\_Externalizing + diagnosis\_cluster ✻  Sex | 0.1667 | 0.01145 | 0.05793 | 0.02036 | 7.468 |
| diagnosis\_cluster + diagnosis\_cluster ✻  Sex | 0.1667 | 0.008112 | 0.04089 | 0.01442 | 7.546 |
| Note.  All models include Sex, Age. | | | | | |

|  |  |  |  |  |  |
| --- | --- | --- | --- | --- | --- |
| 6.3.1 Analysis of Effects - DMPFC | | | | | |
| Effects | P(incl) | P(excl) | P(incl\|data) | P(excl\|data) | BFincl |
| diagnosis\_cluster | 0.6667 | 0.3333 | 0.3138 | 0.6862 | 0.2287 |
| diagnosis\_cluster ✻  Sex | 0.3333 | 0.6667 | 0.01956 | 0.9804 | 0.03991 |
| ASR\_Externalizing | 0.5000 | 0.5000 | 0.7527 | 0.2473 | 3.044 |

|  |  |  |  |  |  |
| --- | --- | --- | --- | --- | --- |
| 6.3.2 Model Averaged Posterior Summary | | | | | |
|  | | | | 95% Credible Interval | |
| Variable | Level | Mean | SD | Lower | Upper |
| Intercept |  | 1.988 | 0.2086 | 1.567 | 2.397 |
| diagnosis\_cluster | 1 | -0.01224 | 0.5128 | -1.048 | 1.007 |
|  | 2 | -0.9011 | 0.4497 | -1.824 | -0.03171 |
|  | 3 | -0.1347 | 0.3335 | -0.8082 | 0.5296 |
|  | 4 | 0.5496 | 0.4641 | -0.3632 | 1.504 |
|  | 6 | -0.1401 | 0.5612 | -1.285 | 0.9634 |
|  | 9 | 0.6385 | 0.3774 | -0.1155 | 1.387 |
| ASR\_Externalizing |  | -0.04996 | 0.02135 | -0.09436 | -0.008804 |
| Sex | f | 0.06156 | 0.1928 | -0.3230 | 0.4459 |
|  | m | -0.06156 | 0.1928 | -0.4510 | 0.3196 |
| Age |  | -0.02252 | 0.01412 | -0.05653 | 0.003826 |
| diagnosis\_cluster ✻  Sex | 1 & f | 0.1933 | 0.4953 | -0.7900 | 1.213 |
|  | 1 & m | -0.1933 | 0.4953 | -1.225 | 0.7770 |
|  | 2 & f | -0.1664 | 0.4047 | -1.006 | 0.6165 |
|  | 2 & m | 0.1664 | 0.4047 | -0.6269 | 0.9958 |
|  | 3 & f | -0.1718 | 0.3127 | -0.8105 | 0.4438 |
|  | 3 & m | 0.1718 | 0.3127 | -0.4516 | 0.7999 |
|  | 4 & f | 0.1413 | 0.4090 | -0.6796 | 0.9658 |
|  | 4 & m | -0.1413 | 0.4090 | -0.9775 | 0.6725 |
|  | 6 & f | 0.3431 | 0.4840 | -0.5811 | 1.365 |
|  | 6 & m | -0.3431 | 0.4840 | -1.380 | 0.5719 |
|  | 9 & f | -0.3394 | 0.3233 | -1.005 | 0.2874 |
|  | 9 & m | 0.3394 | 0.3233 | -0.2932 | 0.9991 |

---

### 6.4 Bayesian ANCOVA Externalising PC

|  |  |  |  |  |  |
| --- | --- | --- | --- | --- | --- |
| Model Comparison | | | | | |
| Models | P(M) | P(M\|data) | BFM | BF10 | error % |
| diagnosis\_cluster | 0.1667 | 0.3967 | 3.288 | 1.000 |  |
| ASR\_Externalizing | 0.1667 | 0.2801 | 1.945 | 0.7060 | 2.786 |
| diagnosis\_cluster + ASR\_Externalizing | 0.1667 | 0.2190 | 1.402 | 0.5521 | 3.188 |
| diagnosis\_cluster + diagnosis\_cluster ✻  Sex | 0.1667 | 0.04220 | 0.2203 | 0.1064 | 3.205 |
| Null model (incl. Sex, Age) | 0.1667 | 0.03984 | 0.2075 | 0.1004 | 2.100 |
| diagnosis\_cluster + ASR\_Externalizing + diagnosis\_cluster ✻  Sex | 0.1667 | 0.02208 | 0.1129 | 0.05566 | 3.430 |
| Note.  All models include Sex, Age. | | | | | |

|  |  |  |  |  |  |
| --- | --- | --- | --- | --- | --- |
| 6.4.1 Analysis of Effects - PC | | | | | |
| Effects | P(incl) | P(excl) | P(incl\|data) | P(excl\|data) | BFincl |
| diagnosis\_cluster | 0.6667 | 0.3333 | 0.6801 | 0.3199 | 1.063 |
| diagnosis\_cluster ✻  Sex | 0.3333 | 0.6667 | 0.06428 | 0.9357 | 0.1374 |
| ASR\_Externalizing | 0.5000 | 0.5000 | 0.5212 | 0.4788 | 1.089 |

|  |  |  |  |  |  |
| --- | --- | --- | --- | --- | --- |
| 6.4.2 Model Averaged Posterior Summary | | | | | |
|  | | | | 95% Credible Interval | |
| Variable | Level | Mean | SD | Lower | Upper |
| Intercept |  | 5.168 | 0.3287 | 4.495 | 5.795 |
| diagnosis\_cluster | 1 | -0.8500 | 0.7158 | -2.301 | 0.5425 |
|  | 2 | -0.2216 | 0.5883 | -1.403 | 0.9358 |
|  | 3 | -0.1166 | 0.4479 | -1.012 | 0.7860 |
|  | 4 | 0.5097 | 0.6344 | -0.7348 | 1.792 |
|  | 6 | -0.8777 | 0.7768 | -2.478 | 0.6104 |
|  | 9 | 1.556 | 0.5168 | 0.5254 | 2.579 |
| ASR\_Externalizing |  | -0.05924 | 0.03305 | -0.1260 | 0.002622 |
| Sex | f | -0.3659 | 0.2652 | -0.8952 | 0.1635 |
|  | m | 0.3659 | 0.2652 | -0.1715 | 0.8917 |
| Age |  | -0.04674 | 0.01917 | -0.09665 | -0.009500 |
| diagnosis\_cluster ✻  Sex | 1 & f | 0.4457 | 0.6993 | -0.9053 | 1.929 |
|  | 1 & m | -0.4457 | 0.6993 | -1.946 | 0.8908 |
|  | 2 & f | -0.3684 | 0.5423 | -1.492 | 0.6827 |
|  | 2 & m | 0.3684 | 0.5423 | -0.6911 | 1.491 |
|  | 3 & f | 0.06996 | 0.4239 | -0.7905 | 0.9087 |
|  | 3 & m | -0.06996 | 0.4239 | -0.9216 | 0.7810 |
|  | 4 & f | 0.2122 | 0.5608 | -0.9180 | 1.353 |
|  | 4 & m | -0.2122 | 0.5608 | -1.364 | 0.9053 |
|  | 6 & f | 0.2379 | 0.6540 | -1.054 | 1.582 |
|  | 6 & m | -0.2379 | 0.6540 | -1.590 | 1.043 |
|  | 9 & f | -0.5974 | 0.4487 | -1.518 | 0.2659 |
|  | 9 & m | 0.5974 | 0.4487 | -0.2759 | 1.504 |

---

### 6.5 Bayesian ANCOVA EXT LTPJ

|  |  |  |  |  |  |
| --- | --- | --- | --- | --- | --- |
| Model Comparison | | | | | |
| Models | P(M) | P(M\|data) | BFM | BF10 | error % |
| diagnosis\_cluster | 0.1667 | 0.3416 | 2.594 | 1.000 |  |
| ASR\_Externalizing | 0.1667 | 0.2584 | 1.742 | 0.7565 | 4.492 |
| diagnosis\_cluster + ASR\_Externalizing | 0.1667 | 0.2075 | 1.309 | 0.6075 | 4.501 |
| Null model (incl. Sex, Age) | 0.1667 | 0.1019 | 0.5675 | 0.2984 | 3.717 |
| diagnosis\_cluster + diagnosis\_cluster ✻  Sex | 0.1667 | 0.05954 | 0.3165 | 0.1743 | 6.672 |
| diagnosis\_cluster + ASR\_Externalizing + diagnosis\_cluster ✻  Sex | 0.1667 | 0.03100 | 0.1600 | 0.09077 | 7.897 |
| Note.  All models include Sex, Age. | | | | | |

|  |  |  |  |  |  |
| --- | --- | --- | --- | --- | --- |
| 6.5.1 Analysis of Effects - LTPJ | | | | | |
| Effects | P(incl) | P(excl) | P(incl\|data) | P(excl\|data) | BFincl |
| diagnosis\_cluster | 0.6667 | 0.3333 | 0.6396 | 0.3604 | 0.8875 |
| diagnosis\_cluster ✻  Sex | 0.3333 | 0.6667 | 0.09054 | 0.9095 | 0.1991 |
| ASR\_Externalizing | 0.5000 | 0.5000 | 0.4969 | 0.5031 | 0.9878 |

|  |  |  |  |  |  |
| --- | --- | --- | --- | --- | --- |
| 6.5.2 Model Averaged Posterior Summary | | | | | |
|  | | | | 95% Credible Interval | |
| Variable | Level | Mean | SD | Lower | Upper |
| Intercept |  | 3.907 | 0.2741 | 3.341 | 4.440 |
| diagnosis\_cluster | 1 | 0.3783 | 0.6795 | -0.9506 | 1.789 |
|  | 2 | -0.9253 | 0.5626 | -2.068 | 0.1762 |
|  | 3 | -0.4716 | 0.4193 | -1.319 | 0.3534 |
|  | 4 | 0.6486 | 0.5870 | -0.5089 | 1.835 |
|  | 6 | -0.6744 | 0.7398 | -2.228 | 0.7411 |
|  | 9 | 1.044 | 0.4664 | 0.1115 | 1.968 |
| ASR\_Externalizing |  | -0.04917 | 0.02864 | -0.1062 | 0.004248 |
| Sex | f | -0.2508 | 0.2573 | -0.7551 | 0.2768 |
|  | m | 0.2508 | 0.2573 | -0.2825 | 0.7488 |
| Age |  | -0.009026 | 0.01777 | -0.05141 | 0.01464 |
| diagnosis\_cluster ✻  Sex | 1 & f | 0.4278 | 0.6739 | -0.8724 | 1.839 |
|  | 1 & m | -0.4278 | 0.6739 | -1.859 | 0.8532 |
|  | 2 & f | -0.2495 | 0.5227 | -1.344 | 0.7525 |
|  | 2 & m | 0.2495 | 0.5227 | -0.7574 | 1.328 |
|  | 3 & f | -0.2119 | 0.4125 | -1.060 | 0.5862 |
|  | 3 & m | 0.2119 | 0.4125 | -0.5973 | 1.055 |
|  | 4 & f | -0.3301 | 0.5461 | -1.465 | 0.7132 |
|  | 4 & m | 0.3301 | 0.5461 | -0.7278 | 1.459 |
|  | 6 & f | 0.9099 | 0.6872 | -0.3552 | 2.385 |
|  | 6 & m | -0.9099 | 0.6872 | -2.395 | 0.3509 |
|  | 9 & f | -0.5462 | 0.4285 | -1.437 | 0.2750 |
|  | 9 & m | 0.5462 | 0.4285 | -0.2866 | 1.429 |

---

### 6.6 Bayesian ANCOVA Externalising RTPJ

|  |  |  |  |  |  |
| --- | --- | --- | --- | --- | --- |
| Model Comparison | | | | | |
| Models | P(M) | P(M\|data) | BFM | BF10 | error % |
| ASR\_Externalizing | 0.1667 | 0.3728 | 2.972 | 1.000 |  |
| Null model (incl. Sex, Age) | 0.1667 | 0.3106 | 2.253 | 0.8332 | 1.930 |
| DiagnoseCluster | 0.1667 | 0.2209 | 1.418 | 0.5927 | 3.986 |
| DiagnoseCluster + ASR\_Externalizing | 0.1667 | 0.08122 | 0.4420 | 0.2179 | 3.481 |
| DiagnoseCluster + DiagnoseCluster ✻  Sex | 0.1667 | 0.01002 | 0.05059 | 0.02687 | 3.401 |
| DiagnoseCluster + ASR\_Externalizing + DiagnoseCluster ✻  Sex | 0.1667 | 0.004453 | 0.02237 | 0.01195 | 25.79 |
| Note.  All models include Sex, Age. | | | | | |

|  |  |  |  |  |  |
| --- | --- | --- | --- | --- | --- |
| 6.6.1 Analysis of Effects - RTPJ | | | | | |
| Effects | P(incl) | P(excl) | P(incl\|data) | P(excl\|data) | BFincl |
| DiagnoseCluster | 0.6667 | 0.3333 | 0.3166 | 0.6834 | 0.2317 |
| DiagnoseCluster ✻  Sex | 0.3333 | 0.6667 | 0.01447 | 0.9855 | 0.02937 |
| ASR\_Externalizing | 0.5000 | 0.5000 | 0.4584 | 0.5416 | 0.8465 |

|  |  |  |  |  |  |
| --- | --- | --- | --- | --- | --- |
| 6.6.2 Model Averaged Posterior Summary | | | | | |
|  | | | | 95% Credible Interval | |
| Variable | Level | Mean | SD | Lower | Upper |
| Intercept |  | 3.637 | 0.2593 | 3.099 | 4.135 |
| DiagnoseCluster | 1 | 0.2251 | 0.6136 | -1.002 | 1.479 |
|  | 2 | -0.5985 | 0.5351 | -1.691 | 0.4468 |
|  | 3 | -0.05632 | 0.3994 | -0.8528 | 0.7416 |
|  | 4 | 0.6734 | 0.5737 | -0.4352 | 1.852 |
|  | 6 | -1.122 | 0.7088 | -2.612 | 0.2094 |
|  | 9 | 0.8786 | 0.4331 | 0.006916 | 1.737 |
| ASR\_Externalizing |  | -0.04271 | 0.02612 | -0.09674 | 0.006600 |
| Sex | f | -0.1196 | 0.2326 | -0.5830 | 0.3379 |
|  | m | 0.1196 | 0.2326 | -0.3431 | 0.5807 |
| Age |  | -0.02180 | 0.01681 | -0.06266 | 0.008912 |
| DiagnoseCluster ✻  Sex | 1 & f | 0.1538 | 0.5916 | -1.024 | 1.365 |
|  | 1 & m | -0.1538 | 0.5916 | -1.376 | 1.006 |
|  | 2 & f | -0.1908 | 0.4682 | -1.154 | 0.7325 |
|  | 2 & m | 0.1908 | 0.4682 | -0.7429 | 1.143 |
|  | 3 & f | 0.1633 | 0.3715 | -0.5759 | 0.9160 |
|  | 3 & m | -0.1633 | 0.3715 | -0.9194 | 0.5706 |
|  | 4 & f | -0.3518 | 0.4939 | -1.391 | 0.6089 |
|  | 4 & m | 0.3518 | 0.4939 | -0.6135 | 1.375 |
|  | 6 & f | 0.1184 | 0.5649 | -1.010 | 1.267 |
|  | 6 & m | -0.1184 | 0.5649 | -1.280 | 0.9952 |
|  | 9 & f | 0.1071 | 0.3820 | -0.6448 | 0.8758 |
|  | 9 & m | -0.1071 | 0.3820 | -0.8827 | 0.6325 |

---

7. # Bayesian ANCOVAs Total Score

### 7.1 Bayesian ANCOVA Total VMPFC

|  |  |  |  |  |  |
| --- | --- | --- | --- | --- | --- |
| Model Comparison | | | | | |
| Models | P(M) | P(M\|data) | BFM | BF10 | error % |
| ASR\_Total | 0.1667 | 0.9266 | 63.11 | 1.000 |  |
| diagnosis\_cluster + ASR\_Total | 0.1667 | 0.06656 | 0.3565 | 0.07184 | 2.478 |
| diagnosis\_cluster + ASR\_Total + diagnosis\_cluster ✻  Sex | 0.1667 | 0.003842 | 0.01929 | 0.004147 | 2.698 |
| diagnosis\_cluster | 0.1667 | 0.001912 | 0.009576 | 0.002063 | 2.521 |
| Null model (incl. Sex, Age) | 0.1667 | 0.001007 | 0.005040 | 0.001087 | 1.510 |
| diagnosis\_cluster + diagnosis\_cluster ✻  Sex | 0.1667 | 8.608×10-5 | 4.304×10-4 | 9.290×10-5 | 5.624 |
| Note.  All models include Sex, Age. | | | | | |

|  |  |  |  |  |  |
| --- | --- | --- | --- | --- | --- |
| 7.1.1 Analysis of Effects - VMPFC | | | | | |
| Effects | P(incl) | P(excl) | P(incl\|data) | P(excl\|data) | BFincl |
| diagnosis\_cluster | 0.6667 | 0.3333 | 0.07240 | 0.9276 | 0.03903 |
| diagnosis\_cluster ✻  Sex | 0.3333 | 0.6667 | 0.003928 | 0.9961 | 0.007888 |
| ASR\_Total | 0.5000 | 0.5000 | 0.9970 | 0.003005 | 331.8 |

|  |  |  |  |  |  |
| --- | --- | --- | --- | --- | --- |
| 7.1.2 Model Averaged Posterior Summary | | | | | |
|  | | | | 95% Credible Interval | |
| Variable | Level | Mean | SD | Lower | Upper |
| Intercept |  | 1.903 | 0.1954 | 1.510 | 2.292 |
| diagnosis\_cluster | 1 | -0.2503 | 0.4932 | -1.265 | 0.7230 |
|  | 2 | -0.2984 | 0.4206 | -1.156 | 0.5235 |
|  | 3 | -0.2978 | 0.3296 | -0.9581 | 0.3530 |
|  | 4 | 0.7000 | 0.4637 | -0.2130 | 1.645 |
|  | 6 | -0.1502 | 0.5482 | -1.276 | 0.9276 |
|  | 9 | 0.2968 | 0.3859 | -0.4683 | 1.077 |
| ASR\_Total |  | -0.02265 | 0.005658 | -0.03566 | -0.01477 |
| Sex | f | -0.07172 | 0.1884 | -0.4539 | 0.3040 |
|  | m | 0.07172 | 0.1884 | -0.3069 | 0.4492 |
| Age |  | -0.01992 | 0.01390 | -0.04931 | 0.001344 |
| diagnosis\_cluster ✻  Sex | 1 & f | -0.06677 | 0.4767 | -1.062 | 0.8763 |
|  | 1 & m | 0.06677 | 0.4767 | -0.8846 | 1.053 |
|  | 2 & f | 0.1783 | 0.3797 | -0.5700 | 0.9551 |
|  | 2 & m | -0.1783 | 0.3797 | -0.9574 | 0.5669 |
|  | 3 & f | -0.02683 | 0.3022 | -0.6400 | 0.5648 |
|  | 3 & m | 0.02683 | 0.3022 | -0.5750 | 0.6380 |
|  | 4 & f | 0.1350 | 0.4010 | -0.6747 | 0.9489 |
|  | 4 & m | -0.1350 | 0.4010 | -0.9610 | 0.6662 |
|  | 6 & f | 0.1341 | 0.4589 | -0.7675 | 1.077 |
|  | 6 & m | -0.1341 | 0.4589 | -1.089 | 0.7544 |
|  | 9 & f | -0.3538 | 0.3195 | -1.014 | 0.2663 |
|  | 9 & m | 0.3538 | 0.3195 | -0.2760 | 1.008 |

---

### 7.2 Bayesian ANCOVA TOTAL MMPFC

|  |  |  |  |  |  |
| --- | --- | --- | --- | --- | --- |
| Model Comparison | | | | | |
| Models | P(M) | P(M\|data) | BFM | BF10 | error % |
| ASR\_Total | 0.1667 | 0.6558 | 9.527 | 1.000 |  |
| diagnosis\_cluster + ASR\_Total | 0.1667 | 0.1501 | 0.8830 | 0.2289 | 3.511 |
| diagnosis\_cluster + ASR\_Total + diagnosis\_cluster ✻  Sex | 0.1667 | 0.1101 | 0.6184 | 0.1678 | 2.855 |
| diagnosis\_cluster | 0.1667 | 0.05386 | 0.2846 | 0.08213 | 6.404 |
| diagnosis\_cluster + diagnosis\_cluster ✻  Sex | 0.1667 | 0.02622 | 0.1346 | 0.03998 | 2.906 |
| Null model (incl. Sex, Age) | 0.1667 | 0.003937 | 0.01976 | 0.006004 | 1.592 |
| Note.  All models include Sex, Age. | | | | | |

|  |  |  |  |  |  |
| --- | --- | --- | --- | --- | --- |
| 7.2.1 Analysis of Effects - MMPFC | | | | | |
| Effects | P(incl) | P(excl) | P(incl\|data) | P(excl\|data) | BFincl |
| diagnosis\_cluster | 0.6667 | 0.3333 | 0.3402 | 0.6598 | 0.2579 |
| diagnosis\_cluster ✻  Sex | 0.3333 | 0.6667 | 0.1363 | 0.8637 | 0.3156 |
| ASR\_Total | 0.5000 | 0.5000 | 0.9160 | 0.08402 | 10.90 |

|  |  |  |  |  |  |
| --- | --- | --- | --- | --- | --- |
| 7.2.2 Model Averaged Posterior Summary | | | | | |
|  | | | | 95% Credible Interval | |
| Variable | Level | Mean | SD | Lower | Upper |
| Intercept |  | 1.443 | 0.2239 | 0.9821 | 1.877 |
| diagnosis\_cluster | 1 | -0.2928 | 0.5668 | -1.440 | 0.8289 |
|  | 2 | -0.1833 | 0.4604 | -1.115 | 0.7316 |
|  | 3 | -0.4437 | 0.3612 | -1.174 | 0.2704 |
|  | 4 | 0.4857 | 0.4961 | -0.4905 | 1.489 |
|  | 6 | -0.4269 | 0.6456 | -1.769 | 0.8135 |
|  | 9 | 0.8609 | 0.4483 | -0.007090 | 1.773 |
| ASR\_Total |  | -0.02023 | 0.006826 | -0.03452 | -0.006068 |
| Sex | f | 0.1022 | 0.2151 | -0.3208 | 0.5371 |
|  | m | -0.1022 | 0.2151 | -0.5413 | 0.3183 |
| Age |  | -0.03074 | 0.01582 | -0.06530 | 4.841×10-4 |
| diagnosis\_cluster ✻  Sex | 1 & f | 0.09266 | 0.5558 | -1.038 | 1.208 |
|  | 1 & m | -0.09266 | 0.5558 | -1.217 | 1.028 |
|  | 2 & f | 0.07521 | 0.4349 | -0.8001 | 0.9387 |
|  | 2 & m | -0.07521 | 0.4349 | -0.9474 | 0.7915 |
|  | 3 & f | 0.1451 | 0.3376 | -0.5339 | 0.8120 |
|  | 3 & m | -0.1451 | 0.3376 | -0.8202 | 0.5214 |
|  | 4 & f | -0.02541 | 0.4553 | -0.9513 | 0.8790 |
|  | 4 & m | 0.02541 | 0.4553 | -0.8878 | 0.9412 |
|  | 6 & f | 0.5708 | 0.5650 | -0.4996 | 1.754 |
|  | 6 & m | -0.5708 | 0.5650 | -1.775 | 0.4904 |
|  | 9 & f | -0.8584 | 0.3734 | -1.620 | -0.1458 |
|  | 9 & m | 0.8584 | 0.3734 | 0.1360 | 1.613 |

---

### 7.3  Bayesian ANCOVA TOTAL DMPFC

|  |  |  |  |  |  |
| --- | --- | --- | --- | --- | --- |
| Model Comparison | | | | | |
| Models | P(M) | P(M\|data) | BFM | BF10 | error % |
| ASR\_Total | 0.1667 | 0.6648 | 9.918 | 1.000 |  |
| diagnosis\_cluster + ASR\_Total | 0.1667 | 0.1747 | 1.058 | 0.2627 | 2.463 |
| diagnosis\_cluster | 0.1667 | 0.07116 | 0.3831 | 0.1070 | 2.366 |
| Null model (incl. Sex, Age) | 0.1667 | 0.06962 | 0.3741 | 0.1047 | 1.677 |
| diagnosis\_cluster + ASR\_Total + diagnosis\_cluster ✻  Sex | 0.1667 | 0.01489 | 0.07557 | 0.02240 | 2.826 |
| diagnosis\_cluster + diagnosis\_cluster ✻  Sex | 0.1667 | 0.004848 | 0.02436 | 0.007292 | 2.415 |
| Note.  All models include Sex, Age. | | | | | |

|  |  |  |  |  |  |
| --- | --- | --- | --- | --- | --- |
| 7.3.1 Analysis of Effects - DMPFC | | | | | |
| Effects | P(incl) | P(excl) | P(incl\|data) | P(excl\|data) | BFincl |
| diagnosis\_cluster | 0.6667 | 0.3333 | 0.2656 | 0.7344 | 0.1808 |
| diagnosis\_cluster ✻  Sex | 0.3333 | 0.6667 | 0.01974 | 0.9803 | 0.04027 |
| ASR\_Total | 0.5000 | 0.5000 | 0.8544 | 0.1456 | 5.867 |

|  |  |  |  |  |  |
| --- | --- | --- | --- | --- | --- |
| 7.3.2 Model Averaged Posterior Summary | | | | | |
|  | | | | 95% Credible Interval | |
| Variable | Level | Mean | SD | Lower | Upper |
| Intercept |  | 2.000 | 0.2068 | 1.577 | 2.398 |
| diagnosis\_cluster | 1 | -0.06637 | 0.5120 | -1.101 | 0.9584 |
|  | 2 | -0.9327 | 0.4520 | -1.857 | -0.04681 |
|  | 3 | -0.06122 | 0.3367 | -0.7336 | 0.6067 |
|  | 4 | 0.5771 | 0.4622 | -0.3454 | 1.506 |
|  | 6 | -0.03459 | 0.5792 | -1.210 | 1.122 |
|  | 9 | 0.5178 | 0.4103 | -0.2980 | 1.328 |
| ASR\_Total |  | -0.01518 | 0.005843 | -0.02799 | -0.006376 |
| Sex | f | 0.1227 | 0.1948 | -0.2685 | 0.5064 |
|  | m | -0.1227 | 0.1948 | -0.5114 | 0.2642 |
| Age |  | -0.02665 | 0.01471 | -0.06486 | 0.003062 |
| diagnosis\_cluster ✻  Sex | 1 & f | 0.1641 | 0.4960 | -0.8241 | 1.189 |
|  | 1 & m | -0.1641 | 0.4960 | -1.194 | 0.8175 |
|  | 2 & f | -0.1798 | 0.4022 | -1.021 | 0.5964 |
|  | 2 & m | 0.1798 | 0.4022 | -0.6034 | 1.015 |
|  | 3 & f | -0.1713 | 0.3177 | -0.8249 | 0.4496 |
|  | 3 & m | 0.1713 | 0.3177 | -0.4540 | 0.8163 |
|  | 4 & f | 0.1683 | 0.4132 | -0.6628 | 0.9993 |
|  | 4 & m | -0.1683 | 0.4132 | -1.010 | 0.6605 |
|  | 6 & f | 0.3857 | 0.4886 | -0.5446 | 1.415 |
|  | 6 & m | -0.3857 | 0.4886 | -1.422 | 0.5366 |
|  | 9 & f | -0.3669 | 0.3315 | -1.047 | 0.2736 |
|  | 9 & m | 0.3669 | 0.3315 | -0.2766 | 1.040 |

---

### 7.4 Bayesian ANCOVA TOTAL PC

|  |  |  |  |  |  |
| --- | --- | --- | --- | --- | --- |
| Model Comparison | | | | | |
| Models | P(M) | P(M\|data) | BFM | BF10 | error % |
| ASR\_Total | 0.1667 | 0.6059 | 7.686 | 1.000 |  |
| diagnosis\_cluster + ASR\_Total | 0.1667 | 0.1922 | 1.189 | 0.3172 | 6.013 |
| diagnosis\_cluster | 0.1667 | 0.1493 | 0.8776 | 0.2464 | 5.424 |
| diagnosis\_cluster + ASR\_Total + diagnosis\_cluster ✻  Sex | 0.1667 | 0.02158 | 0.1103 | 0.03562 | 5.389 |
| diagnosis\_cluster + diagnosis\_cluster ✻  Sex | 0.1667 | 0.01607 | 0.08166 | 0.02652 | 5.550 |
| Null model (incl. Sex, Age) | 0.1667 | 0.01502 | 0.07623 | 0.02479 | 3.764 |
| Note.  All models include Sex, Age. | | | | | |

|  |  |  |  |  |  |
| --- | --- | --- | --- | --- | --- |
| 7.4.1 Analysis of Effects - PC | | | | | |
| Effects | P(incl) | P(excl) | P(incl\|data) | P(excl\|data) | BFincl |
| diagnosis\_cluster | 0.6667 | 0.3333 | 0.3791 | 0.6209 | 0.3053 |
| diagnosis\_cluster ✻  Sex | 0.3333 | 0.6667 | 0.03765 | 0.9624 | 0.07824 |
| ASR\_Total | 0.5000 | 0.5000 | 0.8196 | 0.1804 | 4.543 |

|  |  |  |  |  |  |
| --- | --- | --- | --- | --- | --- |
| 7.4.2 Model Averaged Posterior Summary | | | | | |
|  | | | | 95% Credible Interval | |
| Variable | Level | Mean | SD | Lower | Upper |
| Intercept |  | 5.275 | 0.3153 | 4.606 | 5.862 |
| diagnosis\_cluster | 1 | -0.8718 | 0.7108 | -2.355 | 0.5248 |
|  | 2 | -0.2689 | 0.5883 | -1.447 | 0.9031 |
|  | 3 | -0.05840 | 0.4563 | -0.9793 | 0.8399 |
|  | 4 | 0.5256 | 0.6277 | -0.7223 | 1.786 |
|  | 6 | -0.7178 | 0.8059 | -2.374 | 0.8435 |
|  | 9 | 1.391 | 0.5691 | 0.2486 | 2.504 |
| ASR\_Total |  | -0.02229 | 0.008729 | -0.04341 | -0.008992 |
| Sex | f | -0.2858 | 0.2644 | -0.8194 | 0.2333 |
|  | m | 0.2858 | 0.2644 | -0.2420 | 0.8119 |
| Age |  | -0.05513 | 0.02019 | -0.1070 | -0.02464 |
| diagnosis\_cluster ✻  Sex | 1 & f | 0.4098 | 0.6928 | -0.9318 | 1.876 |
|  | 1 & m | -0.4098 | 0.6928 | -1.881 | 0.9144 |
|  | 2 & f | -0.3664 | 0.5459 | -1.515 | 0.6715 |
|  | 2 & m | 0.3664 | 0.5459 | -0.6872 | 1.505 |
|  | 3 & f | 0.06698 | 0.4193 | -0.7746 | 0.9009 |
|  | 3 & m | -0.06698 | 0.4193 | -0.9083 | 0.7602 |
|  | 4 & f | 0.2362 | 0.5558 | -0.8735 | 1.369 |
|  | 4 & m | -0.2362 | 0.5558 | -1.385 | 0.8624 |
|  | 6 & f | 0.2743 | 0.6515 | -1.029 | 1.606 |
|  | 6 & m | -0.2743 | 0.6515 | -1.628 | 1.007 |
|  | 9 & f | -0.6208 | 0.4474 | -1.549 | 0.2403 |
|  | 9 & m | 0.6208 | 0.4474 | -0.2481 | 1.534 |

---

### 7.5  Bayesian ANCOVA TOTAL RTPJ

|  |  |  |  |  |  |
| --- | --- | --- | --- | --- | --- |
| Model Comparison | | | | | |
| Models | P(M) | P(M\|data) | BFM | BF10 | error % |
| ASR\_Total | 0.1667 | 0.8674 | 32.71 | 1.000 |  |
| diagnosis\_cluster + ASR\_Total | 0.1667 | 0.07312 | 0.3945 | 0.08430 | 7.509 |
| Null model (incl. Sex, Age) | 0.1667 | 0.03266 | 0.1688 | 0.03765 | 5.545 |
| diagnosis\_cluster | 0.1667 | 0.02319 | 0.1187 | 0.02673 | 7.181 |
| diagnosis\_cluster + ASR\_Total + diagnosis\_cluster ✻  Sex | 0.1667 | 0.002583 | 0.01295 | 0.002977 | 7.049 |
| diagnosis\_cluster + diagnosis\_cluster ✻  Sex | 0.1667 | 0.001024 | 0.005123 | 0.001180 | 7.121 |
| Note.  All models include Sex, Age. | | | | | |

|  |  |  |  |  |  |
| --- | --- | --- | --- | --- | --- |
| 7.5.1 Analysis of Effects - RTPJ | | | | | |
| Effects | P(incl) | P(excl) | P(incl\|data) | P(excl\|data) | BFincl |
| diagnosis\_cluster | 0.6667 | 0.3333 | 0.09992 | 0.9001 | 0.05551 |
| diagnosis\_cluster ✻  Sex | 0.3333 | 0.6667 | 0.003606 | 0.9964 | 0.007239 |
| ASR\_Total | 0.5000 | 0.5000 | 0.9431 | 0.05687 | 16.58 |

|  |  |  |  |  |  |
| --- | --- | --- | --- | --- | --- |
| 7.5.2 Model Averaged Posterior Summary | | | | | |
|  | | | | 95% Credible Interval | |
| Variable | Level | Mean | SD | Lower | Upper |
| Intercept |  | 3.688 | 0.2397 | 3.197 | 4.160 |
| diagnosis\_cluster | 1 | 0.2044 | 0.6002 | -0.9918 | 1.410 |
|  | 2 | -0.6744 | 0.5265 | -1.760 | 0.3478 |
|  | 3 | 0.02485 | 0.3978 | -0.7643 | 0.8178 |
|  | 4 | 0.6700 | 0.5553 | -0.4122 | 1.800 |
|  | 6 | -0.7833 | 0.7226 | -2.298 | 0.5957 |
|  | 9 | 0.5584 | 0.4930 | -0.4180 | 1.543 |
| ASR\_Total |  | -0.02045 | 0.006835 | -0.04010 | -0.007296 |
| Sex | f | -0.05322 | 0.2278 | -0.5141 | 0.3988 |
|  | m | 0.05322 | 0.2278 | -0.4028 | 0.5097 |
| Age |  | -0.03221 | 0.01714 | -0.07023 | 0.001866 |
| diagnosis\_cluster ✻  Sex | 1 & f | 0.09076 | 0.5823 | -1.068 | 1.279 |
|  | 1 & m | -0.09076 | 0.5823 | -1.288 | 1.059 |
|  | 2 & f | -0.1746 | 0.4634 | -1.123 | 0.7406 |
|  | 2 & m | 0.1746 | 0.4634 | -0.7572 | 1.117 |
|  | 3 & f | 0.1630 | 0.3690 | -0.5702 | 0.9072 |
|  | 3 & m | -0.1630 | 0.3690 | -0.9173 | 0.5621 |
|  | 4 & f | -0.2925 | 0.4926 | -1.321 | 0.6616 |
|  | 4 & m | 0.2925 | 0.4926 | -0.6746 | 1.310 |
|  | 6 & f | 0.1420 | 0.5605 | -0.9822 | 1.292 |
|  | 6 & m | -0.1420 | 0.5605 | -1.308 | 0.9665 |
|  | 9 & f | 0.07133 | 0.3793 | -0.6882 | 0.8314 |
|  | 9 & m | -0.07133 | 0.3793 | -0.8437 | 0.6811 |

---

### 7.6 Bayesian ANCOVA TOTAL LTPJ

|  |  |  |  |  |  |
| --- | --- | --- | --- | --- | --- |
| Model Comparison | | | | | |
| Models | P(M) | P(M\|data) | BFM | BF10 | error % |
| ASR\_Total | 0.1667 | 0.7035 | 11.86 | 1.000 |  |
| diagnosis\_cluster + ASR\_Total | 0.1667 | 0.1960 | 1.219 | 0.2785 | 7.429 |
| diagnosis\_cluster | 0.1667 | 0.04795 | 0.2518 | 0.06815 | 6.441 |
| diagnosis\_cluster + ASR\_Total + diagnosis\_cluster ✻  Sex | 0.1667 | 0.03154 | 0.1628 | 0.04482 | 3.758 |
| Null model (incl. Sex, Age) | 0.1667 | 0.01321 | 0.06692 | 0.01877 | 2.735 |
| diagnosis\_cluster + diagnosis\_cluster ✻  Sex | 0.1667 | 0.007831 | 0.03946 | 0.01113 | 4.907 |
| Note.  All models include Sex, Age. | | | | | |

|  |  |  |  |  |  |
| --- | --- | --- | --- | --- | --- |
| 7.6.1 Analysis of Effects - LTPJ | | | | | |
| Effects | P(incl) | P(excl) | P(incl\|data) | P(excl\|data) | BFincl |
| diagnosis\_cluster | 0.6667 | 0.3333 | 0.2833 | 0.7167 | 0.1976 |
| diagnosis\_cluster ✻  Sex | 0.3333 | 0.6667 | 0.03937 | 0.9606 | 0.08196 |
| ASR\_Total | 0.5000 | 0.5000 | 0.9310 | 0.06899 | 13.50 |

|  |  |  |  |  |  |
| --- | --- | --- | --- | --- | --- |
| 7.6.2 Model Averaged Posterior Summary | | | | | |
|  | | | | 95% Credible Interval | |
| Variable | Level | Mean | SD | Lower | Upper |
| Intercept |  | 3.958 | 0.2572 | 3.432 | 4.467 |
| diagnosis\_cluster | 1 | 0.3320 | 0.6503 | -0.9348 | 1.691 |
|  | 2 | -1.003 | 0.5595 | -2.155 | 0.07703 |
|  | 3 | -0.3704 | 0.4215 | -1.223 | 0.4655 |
|  | 4 | 0.6651 | 0.5766 | -0.4743 | 1.840 |
|  | 6 | -0.3238 | 0.7547 | -1.889 | 1.130 |
|  | 9 | 0.7003 | 0.5166 | -0.3096 | 1.741 |
| ASR\_Total |  | -0.02244 | 0.007528 | -0.04262 | -0.01050 |
| Sex | f | -0.1844 | 0.2454 | -0.6745 | 0.3079 |
|  | m | 0.1844 | 0.2454 | -0.3182 | 0.6694 |
| Age |  | -0.02077 | 0.01845 | -0.06996 | 0.009785 |
| diagnosis\_cluster ✻  Sex | 1 & f | 0.3219 | 0.6670 | -0.9780 | 1.696 |
|  | 1 & m | -0.3219 | 0.6670 | -1.718 | 0.9647 |
|  | 2 & f | -0.2229 | 0.5228 | -1.294 | 0.7827 |
|  | 2 & m | 0.2229 | 0.5228 | -0.7934 | 1.282 |
|  | 3 & f | -0.2059 | 0.4029 | -1.024 | 0.5820 |
|  | 3 & m | 0.2059 | 0.4029 | -0.5907 | 1.021 |
|  | 4 & f | -0.2555 | 0.5385 | -1.371 | 0.7798 |
|  | 4 & m | 0.2555 | 0.5385 | -0.7964 | 1.368 |
|  | 6 & f | 0.9402 | 0.6900 | -0.3349 | 2.447 |
|  | 6 & m | -0.9402 | 0.6900 | -2.453 | 0.3192 |
|  | 9 & f | -0.5778 | 0.4260 | -1.454 | 0.2462 |
|  | 9 & m | 0.5778 | 0.4260 | -0.2573 | 1.448 |
